# Resurrection of the Plant Immune Receptor Sr50 to Overcome Pathogen Immune Evasion

**DOI:** 10.1101/2024.08.07.607039

**Authors:** Kyungyong Seong, Wei Wei, Sophie C Sent, Brandon Vega, Amanda Dee, Griselda Ramirez-Bernardino, Rakesh Kumar, Lorena Parra, Isabel ML Saur, Ksenia Krasileva

## Abstract

Pathogen-driven plant diseases cause significant crop losses worldwide. The introgression of intracellular nucleotide-binding leucine-rich repeat receptor (NLR) genes into elite crop cultivars is a common strategy for disease control, yet pathogens rapidly evolve to evade NLR-mediated immunity. The NLR gene *Sr50* protects wheat against stem rust, a devastating disease caused by the fungal pathogen *Puccinia graminis* f. sp. *tritici* (*Pgt*). However, mutations in AvrSr50 allowed *Pgt* to evade Sr50 recognition, leading to resistance breakdown. Advances in protein structure modeling can enable targeted NLR engineering to restore recognition of escaped effectors. Here, we combined iterative computational structural analyses and site-directed mutagenesis to engineer Sr50 recognition of AvrSr50^QCMJC^, a *Pgt* effector variant that evades wild-type Sr50 detection. Derived by molecular docking, our initial structural model identified the K711D substitution in Sr50, which partially restored AvrSr50^QCMJC^ recognition. Enhancing Sr50^K711D^ expression via strong promoters compensated for weak recognition and restored robust immune responses. Further structural refinements led to the generation of five double and two triple receptor mutants. These engineered mutants, absent in nature, showed robust dual recognition for AvrSr50 and AvrSr50^QCMJC^ in both *Nicotiana benthamiana* and wheat protoplasts. Notably, the K711D substitution was essential and synergistic with the additional substitutions for AvrSr50^QCMJC^ recognition, demonstrating protein epistasis. Furthermore, this single substitution altered AlphaFold 2 predictions, enabling accurate modeling of the Sr50^K711D^–AvrSr50 complex structure, consistent with our final structural hypothesis. Collectively, this study outlines a framework for NLR engineering to counteract pathogen adaptation and provides novel Sr50 variants with potential for stem rust resistance.

## Introduction

Intracellular nucleotide-binding leucine-rich repeat receptors (NLRs) are crucial components of plant immunity, functioning as molecular sentinels that detect and respond to pathogen threats and thereby mediating disease resistance (Jones and Dangl, 2006). Upon recognizing pathogen effectors, NLRs trigger immune responses, often culminating in localized cell death termed the hypersensitive response (HR) (Dodds and Rathjen, 2010). These immune responses can significantly restrict pathogen proliferation, making NLRs vital targets for crop genetic protection strategies (Arora *et al*., 2019; Dangl *et al*., 2013).

Among many NLRs deployed in breeding, *Sr50* has been introgressed from rye into bread wheat to confer protection against devastating stem rust disease caused by the fungal pathogen *Puccinia graminis* f. sp. *tritici* (*Pgt*) (Mago *et al*., 2015). Sr50’s C-terminal leucine-rich repeat (LRR) domain directly binds the *Pgt* effector AvrSr50, leading to resistance to *Pgt* (Tamborski *et al*., 2023). However, mutations in AvrSr50 allow *Pgt* to evade Sr50-mediated recognition, compromising immunity (Möller and Stukenbrock, 2017; Sánchez-Vallet *et al*., 2018). Specifically, the AvrSr50^QCMJC^ variant secreted by *Pgt* isolate QCMJC turns stealthy with the Q121K mutation on the protein surface (Chen *et al*., 2017; Ortiz *et al*., 2022). Such rapid effector evolution underlying NLR recognition escape poses a challenge to crop protection strategies and necessitates innovative approaches to restore and enhance NLR-mediated resistance (Märkle *et al*., 2022).

Addressing recognition evasion in a rapid and rational manner requires engineering approaches, yet progress has been hindered by the vast mutational space in NLRs and limited understanding of NLR-effector interactions (Tamborski *et al*., 2023; Zdrzałek *et al*., 2023). For example, varying 12 surface-exposed residues constituting a typical effector binding site would result in up to 20^12^ NLR allelic variants (Tamborski *et al*., 2023). Prioritization and rational selection of compensatory mutations needed to restore effector recognition by NLRs require development of new structure-informed computational strategies. Structural elucidation techniques, such as cryogenic electron microscopy (Cryo-EM), can aid development of design strategies. However, experimental NLR structure determination methods are challenging and resource-intensive with only a limited number of NLR-effector complex structures having been resolved (Förderer *et al*., 2022; Lawson *et al*., 2024; Ma *et al*., 2020; Martin *et al*., 2020; Zhao *et al*., 2022). Computational approaches, such as AlphaFold 2 (AF2) and AF3, offer promising alternatives by predicting NLR-effector complex structures (Abramson *et al*., 2024; Evans *et al*., 2021; Jumper *et al*., 2021), but their accuracy are often unpredictable using the standard metrics and has been challenged by discrepancies with recently solved Cryo-EM structure of MLA13 and its effector AVR_A13_-1 (Lawson *et al*., 2024).

Despite inherent difficulties, NLR engineering has significantly progressed over the past few years. Most successful examples utilize small integrated domains found in a subset of NLRs as platforms for effector binding (Baggs *et al*., 2017; Bentham *et al*., 2023; Cesari *et al*., 2022; De La Concepcion *et al*., 2019; Kroj *et al*., 2016; Maidment *et al*., 2023; Sarris *et al*., 2016). In contrast, engineering the C-terminal LRR domain remains a challenging pursuit, although it is postulated to mediate effector binding across many NLRs. Most efforts resort to gain-of-function random mutagenesis to counter escape effector mutants (Farnham and Baulcombe, 2006; Harris *et al*., 2013; Huang *et al*., 2021; Segretin *et al*., 2014). We recently leveraged the natural NLR sequence diversity, pinpointing and targeting rapidly evolving, highly variable (hv) residues to switch recognition specificity between two related NLRs (Prigozhin and Krasileva, 2021; Tamborski *et al*., 2023). However, utilizing natural diversity data from receptors alone does consider their structural interplay with effectors. When effector genes mutate, numerous receptor residues need to be re-screened, making it difficult to efficiently restore NLR recognition along pathogen evolution. Therefore, there is a pressing need for methodologies that can incorporate the conformation of NLR-effector complexes for prioritization of targeted mutations.

In this study, we developed a computationally informed rational NLR improvement approach and engineered Sr50 to resurrect its function against the escape mutant AvrSr50^QCMJC^. Using structural analyses combined with molecular docking and site-directed mutagenesis, we iteratively inferred, tested, and refined the Sr50 and AvrSr50 interaction models. Our approach identified a key mutation, K711D, that weakly restored AvrSr50^QCMJC^ recognition. Interestingly, this substitution functioned synergistically with other additional mutations and was essential for double and triple mutants to elicit robust HR against AvrSr50^QCMJC^ in *Nicotiana benthamiana*. The engineered mutants consistently demonstrated recognition in wheat protoplasts, confirming successful engineering. Furthermore, the K711D substitution enabled AF2-Multimer to successfully predict the Sr50^K711D^–AvrSr50 complex, even though it failed to model the wild type Sr50–AvrSr50 complex. Our study not only provides key insights into Sr50–AvrSr50 interactions but also establishes a framework for rational engineering of LRRs to counteract effector evolution. Engineered Sr50 variants with expanded recognition specificity can mediate stem rust resistance caused by a large set of *Pgt* isolates present in the global pathogen population.

## Results

### Conceptual framework for the iterative refinement of structural hypotheses

The goal of this study is to develop a reliable structural model of Sr50 and AvrSr50 interactions through an iterative approach, which can subsequently offer engineering strategies for AvrSr50^QCMJC^ recognition. To achieve this, we built a conceptual framework based on iterative refinement of structural models (Fig. 1a). Initially, we generated multiple initial structural hypotheses with molecular docking simulations. Based on these models, we prioritized introducing mutations primarily at charged or polar residues in Sr50 and AvrSr50, hypothesizing that these residues would influence ligand-receptor specificity. Each mutation was designed to either disrupt effector recognition or mediate recognition escape, resulting in the loss of HR. However, complementary mutations in neighboring residues of the interacting partner may restore the immune phenotype. This interplay between mutations provides evidence for residual proximity for Sr50 and AvrSr50, which we subsequently leveraged to refine our structural hypotheses. This iterative process progressively incorporates residue constraints, leading to a more reliable model that guides the introduction of mutations enabling Sr50 to gain AvrSr50^QCMJC^ recognition.

**Figure 1.**
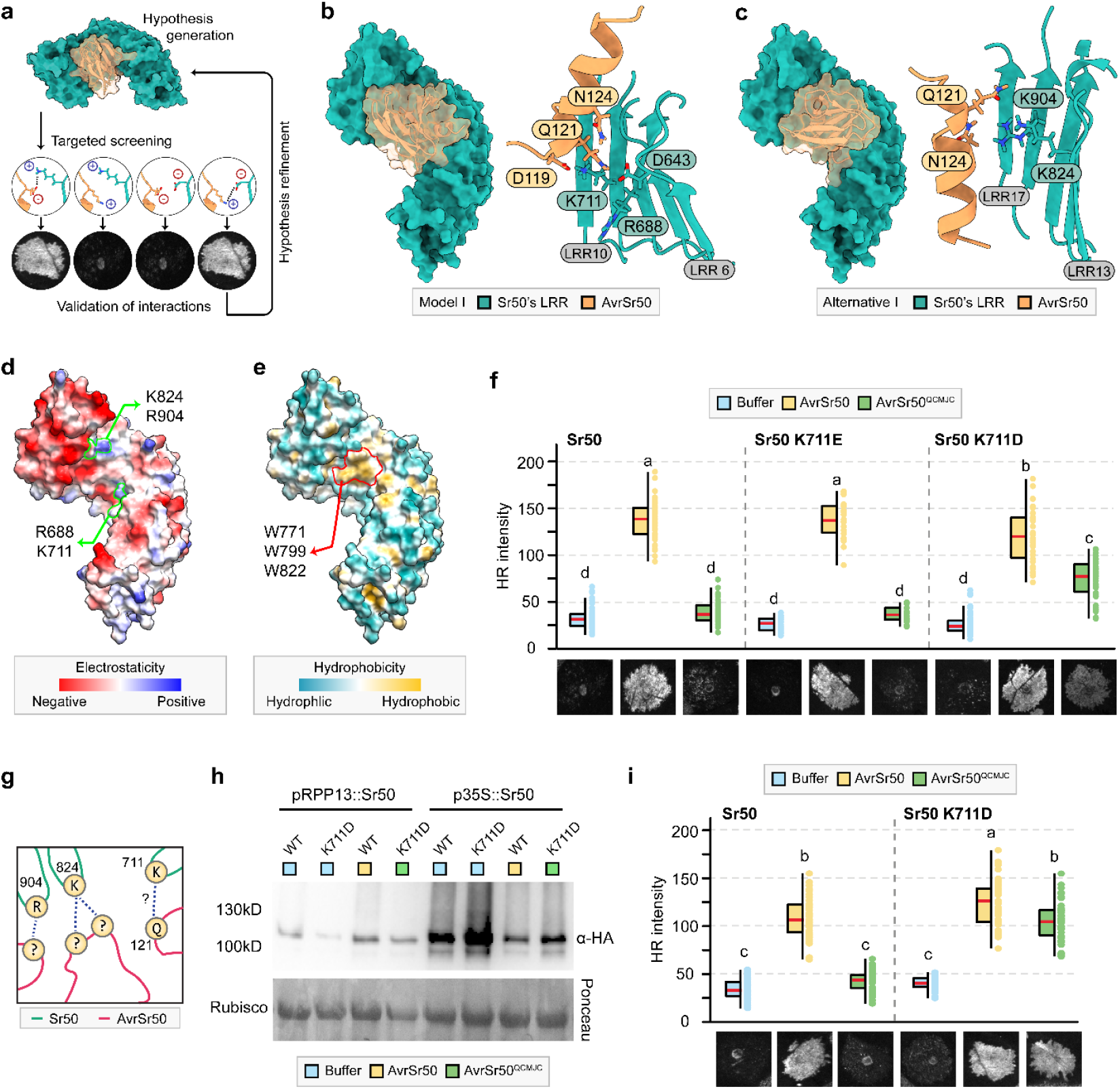
Initial structural hypotheses derived from molecular docking simulations enable the gain of AvrSr50^QCMJC^ recognition. **(a)** Conceptual framework for the iterative refinement of structural hypotheses. **(b-c)** Predicted poses of Sr50 and AvrSr50 through molecular docking simulations. Some parts of a loop between β2–β3, mostly including an unstructured region, are removed from AvrSr50 for visualization (positions 42-66). The local environment around AvrSr50’s Q121 is visualized. **(d and e)** Estimated electrostatic potential and hydrophobicity of the leucine-rich repeat (LRR) domain in Sr50. **(f and i)** Cell death phenotypes on *Nicotiana benthamiana*. Indicated pairs of receptors and effectors were co-infiltrated, and the hypersensitive response (HR) intensity was quantified at three days post infiltration. The statistical significance was accessed with one-way ANOVA followed by a post-hoc Tukey Honestly Significant Difference test. Groups sharing the same letter are not significantly different (*P* > 0.05). Representative HR images corresponding to the median intensity are displayed. **(f)** Receptor expressions were driven by pRPP13. **(i)** Receptor expressions were driven by p35S. **(g)** Simplified schematic representation of the proposed interaction interface between Sr50 and AvrSr50. Amino acids are shown in circles, with their positions labeled adjacent to each residue for Sr50 on green lines and AvrSr50 on red lines. Blue dashed lines indicate proposed interacting pairs. **(h)** Representative Western blot for the wild type Sr50 (WT) and the Sr50^K711D^ mutant under pRPP13 or p35S upon expression in *N. benthamiana*.

### Initial structural hypothesis generation with molecular docking

To formulate an initial structural hypothesis, we modeled the Sr50 and AvrSr50 complex with AF2-Multimer and AF3, but these predictions showed low accuracy (Fig. S1). Thus, we used molecular docking algorithms guided by structural criteria drawn from the Sr35 resistosome and the assumption that AvrSr50’s Q121 directly interacts with Sr50’s LRR residues (Förderer *et al*., 2022; Ortiz *et al*., 2022; Zhao *et al*., 2022). Our intention was to diversify initial models and identify experimentally testable hypotheses.

Docking simulations produced three distinct models (Fig. 1b-c; Fig. S2). We prioritized two that aligned with our assumption: the Q121K substitution in AvrSr50^QCMJC^ creates repulsion with Sr50’s positively charged residues, impeding recognition. In Model I (Fig. 1b), AvrSr50’s Q121 was proximal to Sr50’s arginine and lysine at positions 688 and 711. An alternative model positioned Q121 close to lysine and arginine at positions 824 and 904 (Fig. 1c). We hypothesized that these four charged residues may determine effector specificity (Fig. 1d), while a neighboring cluster of tryptophan residues across LRRs 11 to 13 influence binding affinity (Fig. 1e).

### Sr50^K711D^ gains AvrSr50^QCMJC^ recognition without losing AvrSr50 recognition

We introduced aspartic and glutamic acid substitutions into R688, K711, K824 or R904 in Sr50, aiming to stabilize repulsion from Q121K of AvrSr50^QCMJC^. Co-infiltrating the single mutants with AvrSr50 and AvrSr50^QCMJC^ in *N. benthamiana* revealed the Sr50^K824D/E^ and Sr50^R904D/E^ mutants lost AvrSr50-dependent HR (Fig. S3), potentially indicating their involvement in effector recognition (Fig. 1g). However, they could not induce HR against AvrSr50^QCMJC^ (Fig. S3). Similarly, the Sr50^R688D/E^ mutants failed to gain AvrSr50^QCMJC^ recognition (Fig. S3). Sr50^K711D^ and Sr50^K711E^ retained AvrSr50 recognition, but only Sr50^K711D^ induced AvrSr50^QCMJC^-dependent cell death (Fig. 1f; Table S1). Their HR was weaker compared to Sr50 and AvrSr50, suggesting a suboptimal interaction. This weak interaction would make Sr50^K711D^-mediated resistance susceptible to being overcome by the rapid evolution of AvrSr50^QCMJC^ upon employment of Sr50^K711D^ in the field. Nevertheless, these results indicated that Sr50^K711D^ gained recognition against AvrSr50^QCMJC^ while maintaining its interaction with AvrSr50 (Fig. 1g).

### Enhanced expression compensates for weak recognition of AvrSr50^QCMJC^ by Sr50^K711D^

To determine whether the increased HR of Sr50^K711D^ against AvrSr50^QCMJC^ was influenced by protein abundance, we performed a western blot analysis of *N. benthamiana* transiently expressing the constructs (Fig. 1h). Sr50^K711D^ consistently showed lower protein levels compared to Sr50 under the RPP13 promoter, indicating that its HR against the escape mutant was not due to elevated protein abundance. When expressed under the strong constitutive 35S promoter, the protein levels of Sr50^K711D^ exceeded those of Sr50 and those under the RPP13 promoter (Fig. 1h). Consistently, Sr50^K711D^ showed enhanced HR for both AvrSr50 and AvrSr50^QCMJC^ (Fig. 1i), whereas Sr50 remained unresponsive to AvrSr50^QCMJC^. This indicated that elevated expression driven by the 35S promoter compensated for weak recognition of AvrSr50^QCMJC^ by Sr50^K711D^ for enhanced immune responses. Although promoter engineering is a viable engineering strategy, we continued our pursuit with protein structure-guided engineering driven by the RPP13 promoter to further optimize Sr50^K711D^ recognition through additional amino acid mutations.

### Sr50^K711D^ does not recognize the AvrSr50^Q121K^ single mutant

The gain of function in Sr50^K711D^ was informed by but does not serve as direct validation of the existing structural hypothesis (Fig. 1b). The mature forms of AvrSr50 and AvrSr50^QCMJC^ have ten substitutions, including Q121K, which directly mediates recognition escape (Fig. 2a; Fig. S4) (Ortiz *et al*., 2022). It was therefore crucial to determine whether Sr50^K711D^ could induce HR against the AvrSr50^Q121K^ single mutant. Consistent with the previous study (Ortiz *et al*., 2022), reverting K121 of AvrSr50^QCMJC^ to glutamine was sufficient to restore the HR for both Sr50 and Sr50^K711D^ (Fig. 2b; Table S1). However, Sr50^K711D^ did not trigger HR against AvrSr50^Q121K^. This result suggested that additional effector residues may contribute to the interactions between Sr50’s K711 and AvrSr50’s Q121 (Fig. 2c) or that they do not interact with each other (Fig. 2d).

**Figure 2.**
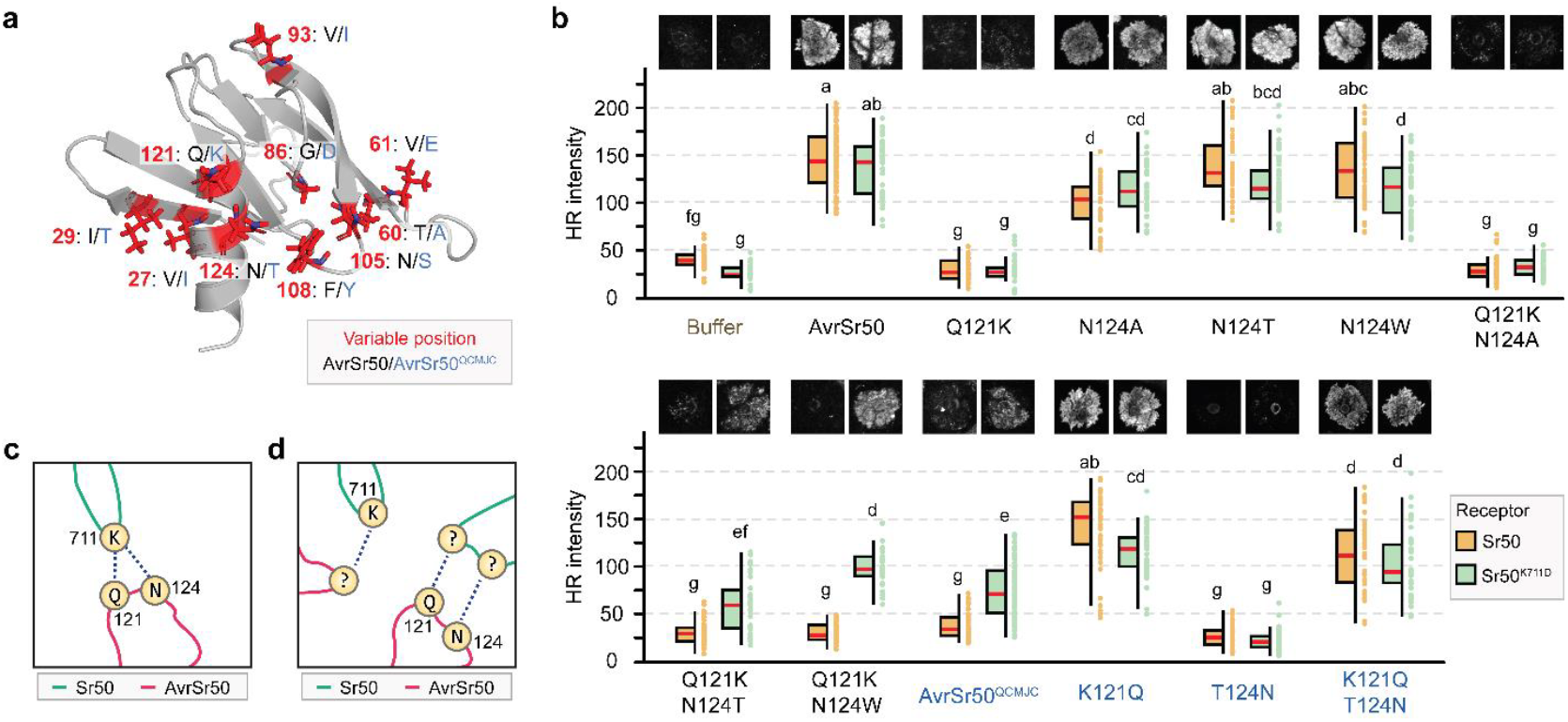
AvrSr50’s Q121 and N124 interact with other residues than K711 in Sr50. **(a)** Amino acid differences between AvrSr50 and AvrSr50^QCMJC^. The two effectors differ by 10 residues in their mature forms, with differences shown in black for AvrSr50 and blue for AvrSr50^QCMJC^. **(b)** Cell death phenotypes on *Nicotiana benthamiana*. Sr50 (orange) or Sr50^K711D^ (green) expressed under pRPP13 was co-infiltrated with the indicated effector variant. These include buffer, AvrSr50 variants (black) and AvrSr50^QCMJC^ variants (blue). The HR intensity was quantified at three days post infiltration. The statistical significance was accessed with one-way ANOVA followed by a post-hoc Tukey Honestly Significant Difference test. Groups sharing the same letter are not significantly different (*P* > 0.05). Representative HR images corresponding to the median intensity are displayed. **(c** and **d)** Simplified schematic representation of the proposed interaction interface between Sr50 and AvrSr50. Amino acids are shown in circles, with their positions labeled adjacent to each residue for Sr50 on green lines and AvrSr50 on red lines. Blue dashed lines indicate proposed interacting pairs.

### Sr50’s K711 does not directly interact with AvrSr50’s Q121

AvrSr50^QCMJC^ carries the N124T substitution adjacent to Q121K along the terminal α-helix (Fig. 2a). We examined whether this additional mutation contributes to the differential responses of Sr50^K711D^ towards AvrSr50^QCMJC^ and AvrSr50^Q121K^ (Fig. 2c). Experiments showed that the AvrSr50^N124T^ single mutant did not significantly impact the activation of Sr50 and Sr50^K711D^, although it slightly attenuated the HR responses compared to AvrSr50 (Fig. 2b). In contrast, co-infiltration of Sr50^K711D^ and the AvrSr50^Q121K/N124T^ double mutant partially restored the cell death phenotype. Also, AvrSr50^QCMJC K121Q/T124N^ tended to reduce the intensity of HR, compared to AvrSr50^QCMJC K121Q^, suggesting that position 124 contributes to the interaction but is not a major determinant.

To further explore the role of this position and distinguish between alternative explanations (Fig. 2c-d), we substituted AvrSr50’s N124 with two structurally distinct amino acids: alanine and tryptophan. We initially hypothesized that alanine, due to its similar size to threonine, would enhance immune activation of Sr50^K711D^ when introduced into the AvrSr50^Q121K^ background, similar to AvrSr50^Q121K/N124T^. In contrast, we predicted that the bulky, highly hydrophobic tryptophan would have the opposite effect, completely abolishing the interaction with Sr50^K711D^. Unexpectedly, AvrSr50^N124A^ diminished cell death responses for both Sr50 and Sr50^K711D^, and AvrSr50^Q121K/N124A^ failed to enhance HR (Fig. 2b). In contrast, the AvrSr50^Q121K/N124W^ double mutant increased HR when co-infiltrated with Sr50^K711D^, compared to AvrSr50^Q121K^ (Fig. 2b). However, this enhanced HR was also observed in the co-infiltration with Sr50^K711A^ and Sr50^K711E^ (Fig. S5). This lack of specific association with the K711D substitution suggested that AvrSr50’s Q121K and N124T may interact with other receptor residues instead of Sr50’s K711 (Fig. 2d).

### Sr50’s K824 and R904 form an additional contact with AvrSr50

Given the analysis of the additional mutants, we concluded that our initial docking model was not fully accurate, and therefore we decided to explore additional receptor-effector contact points. As mutating Sr50’s K824 or R904 to aspartic and glutamic acids abolished or attenuated AvrSr50 recognition (Fig. 1g; Fig. S3), we hypothesized that K824 and R904 interact with negatively charged effector residues. Mapping these residues in Model I identified two aspartic acids and three glutamic acids (Fig. 1b and 3a), forming a negatively charged surface potentially interfacing the LRR domain (Fig. 3b). We prioritized AvrSr50’s D30, E115 and E117, excluding E35, possibly stabilized by K28 and K79 by internal interactions (Fig. S6), and D119, positioned near Q121 and unlikely to contact K824 or R904 in Model I (Fig. 1b).

**Figure 3.**
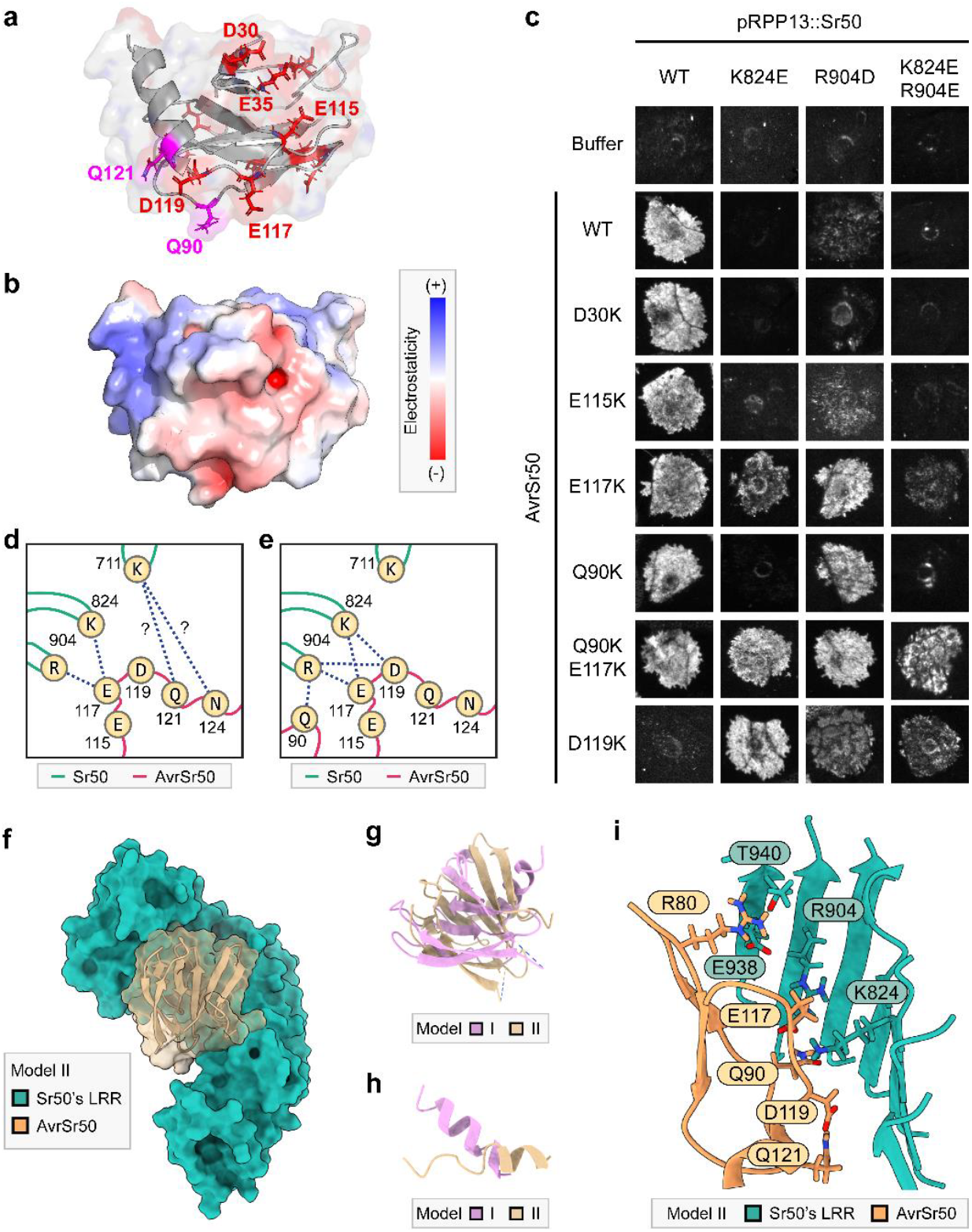
Sr50’s terminal leucine-rich repeats form a critical effector binding interface. **(a** and **b)** Predicted structure of AvrSr50 in the identical orientation. **(a)** Negatively charged amino acids are indicated in red with Q90 and Q121 highlighted in magenta. **(b)** Estimated electrostatic potential is mapped to the surface of AvrSr50. **(c)** Cell death phenotypes on *Nicotiana benthamiana*. Indicated receptor and effector pairs were co-infiltrated, and the phenotypes were recorded at three days post infiltration. Representative HR images corresponding to the median intensity are displayed. **(d** and **e)** Simplified schematic representation of the proposed interaction interface between Sr50 and AvrSr50. Amino acids are shown in circles, with their positions labeled adjacent to each residue for Sr50 on green lines and AvrSr50 on red lines. Blue dashed lines indicate proposed interacting pairs. **(f)** Predicted poses of Sr50 and AvrSr50 in Model II. Some parts of a loop between β2–β3, mostly including an unstructured region, are removed from AvrSr50 for visualization (positions 42-66). **(g** and **h)** Superposition of AvrSr50 in Model I and II. The entire complex structures of the two models were superposed to fix the coordinates of Sr50 consistent. The AvrSr50 structures are then visualized without positions 42 to 66. **(h)** Only the terminal α-helix of AvrSr50 is displayed from the superposed AvrSr50 structures. **(i)** Local environment around Sr50’s K824 and R904 in Model II.

We created AvrSr50^D30K^, AvrSr50^E115K^ and AvrSr50^E117K^ and co-infiltrated them with Sr50^K824E^ or Sr50^R904D^ to test for HR restoration (Fig. 3c; Fig. S7; Table S1). AvrSr50^D30K^ had no notable effects on cell death induction. AvrSr50^E115K^ induced weak HR against Sr50^R904D^, only slightly stronger than Sr50^R904D^ and AvrSr50. Notably, AvrSr50^E117K^ induced strong cell death against both Sr50^K824E^ and Sr50^R904D^, indicating that E117 interacts proximally with Sr50’s K824 and R904.

### Refining structural hypotheses with experimental constraints

We refined our structural hypothesis, using ColabDock with four residue pairs from Sr50 and AvrSr50 as constraints: experimentally derived K824-E117 and R904-E117, and postulated K711-Q121 and K711-N124 from Model I (Fig. 3d). The updated Model II showed notable changes in AvrSr50’s positioning compared to Model I (Fig. 3f and 3g). However, AvrSr50’s terminal α-helix structure was distorted to accommodate the specified restraints (Fig. 3h). Close inspection revealed that the constraints from Model I were likely incorrect, as our experimental outcomes indicated (Fig. 2d). Specifically, the distance between AvrSr50’s E117 and Q121 was comparatively smaller than the distance between Sr50’s K711 and K824 (Fig. 1d; Fig. 3a), making the K711-Q121 and K711-N124 interactions impossible, when the K824-E117 and R904-E117 constraints were satisfied. However, removing these K711 restraints resulted in prediction models where no AvrSr50 residues were positioned near Sr50’s K711 (Fig. S8), not aligning with our postulation that K711 is involved in the interaction (Fig. 2d). We proceeded with Model II as our next structural hypothesis while acknowledging its limitation.

### Additional AvrSr50 residues contact Sr50’s K824 and R904

Model II revealed additional charged or polar effector residues which may be proximal to Sr50’s K824 and R904: Q90 and D119 (Fig. 3a and 3i). Unlike in Model I, AvrSr50’ D119 in Model II lied much closer to these receptor residues. We tested whether AvrSr50^Q90K^ and AvrSr50^D119K^ could restore the HR for Sr50^K824E^ and Sr50^R904D^ (Fig. 3c; Fig. S7; Table S1). AvrSr50^Q90K^ could induce cell death via Sr50^R904D^ but not Sr50^K824E^. Furthermore, the AvrSr50^Q90K/E117K^ double mutant triggered robust HR with the Sr50^K824E/R904E^ double mutant, a response absent in tested single effector mutants, supporting the interaction between AvrSr50’s Q90 and Sr50’s R904 (Fig. 3e). Similarly, AvrSr50^D119K^ strongly restored recognition by Sr50^K824E^ and induced comparatively weak HR with Sr50^R904D^ and Sr50^K824E/R904E^ (Fig. 3c; Fig. S7). Notably, AvrSr50^D119K^ abolished interaction with Sr50, suggesting D119 may be critical for recognition, interacting with Sr50’s K824 and positioned near R904. Collectively, we could identify five residue pairs as key components of this interaction interface (Fig. 3e).

### Positively charged residues on the terminal LRR lead to auto-activity

Following Model II, we attempted testing the relevance of the interaction between Sr50’s E938 and AvrSr50’s R80 *in planta* (Fig. 3i). However, Sr50^E938K^ showed strong auto-activity, making it challenging to discern phenotypic changes upon co-infiltration with effector mutants (Fig. S9). Alternatively, we expressed Sr50^T940K^ and Sr50^T940R^ mutants, but they also triggered severe auto-activity (Fig. S9). These findings suggested that the very terminal LRR unit might play a role in stabilizing inactive Sr50, and introducing positively charged residues at these positions disrupts Sr50’s stability.

### AvrSr50’s α-helix enriched with positively charged residues contact the inner concave of the LRR domain

Model II had implausible structural features, such as R128 clashing into the receptor backbones (Fig. 3h and 4a). Despite these limitations, we noted potential electrostatic complementarity in this region. Negatively charged amino acids, D589, D618, D641 and D643, are proximal to Sr50’s K711 (Fig. 4a). Although some of these residues contact the NB-ARC domain potentially for interdomain stabilization and may not be available for interactions (Fig. 4b), the others could participate in the interaction with the effector. Notably, AvrSr50 contains positively charged H125, R128, R129, H131 and R132 at the end of the terminal α-helix (Fig. 4c). This cluster creates strong positive electrostatic potential (Fig. 4d) and is potentially positioned near the negatively charged amino acids of Sr50 in Model II (Fig. 4a). Among these residues, R128 lies in the same plane with E117, D119, Q121 and N124 which were shown to alter the interaction with Sr50 mutants (Fig. 4c).

**Figure 4.**
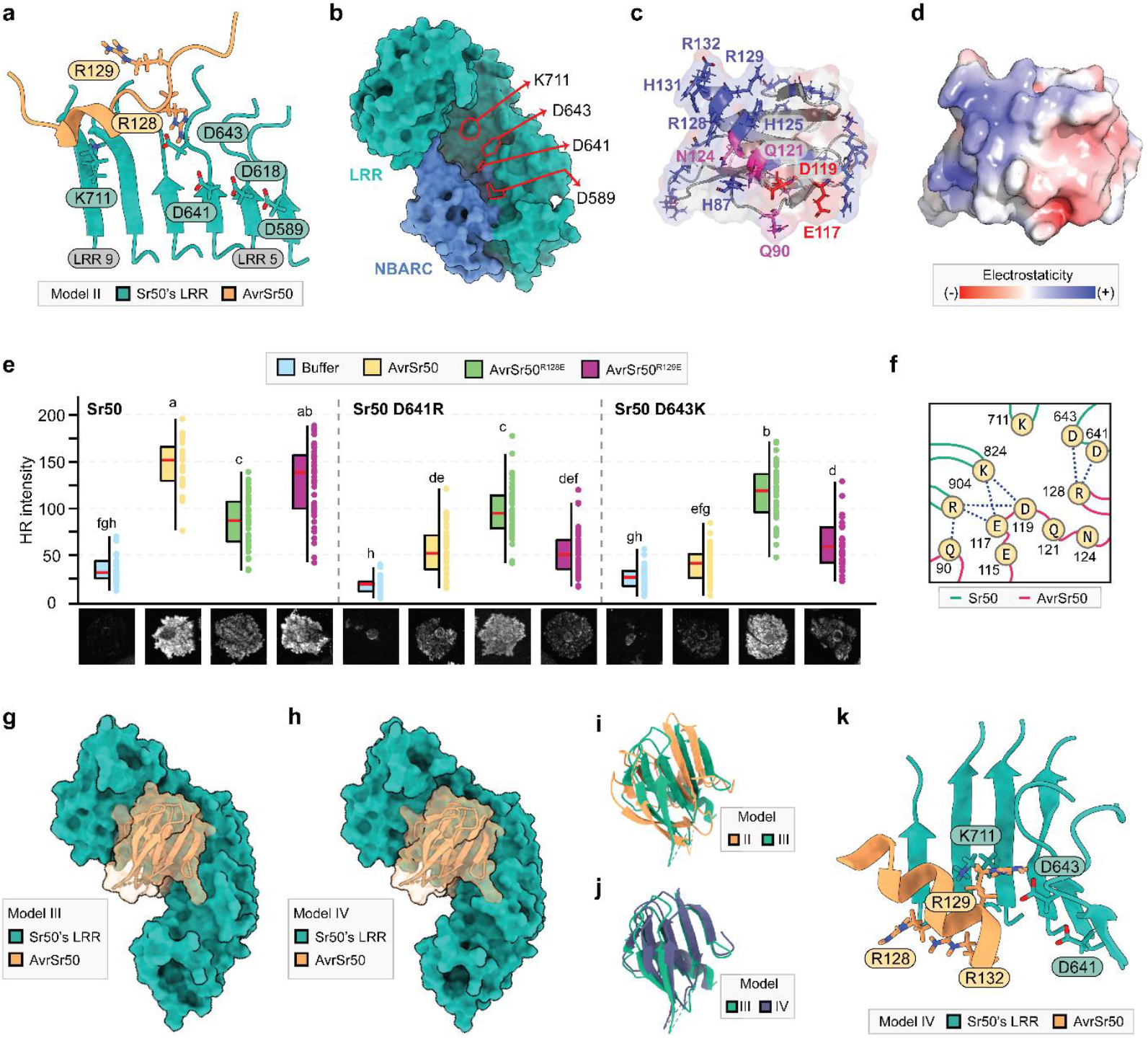
AvrSr50’s α-helix enriched with positively charged residues contact the inner concave of the LRR domain. **(a)** Local environment around Sr50’s K711 in Model II. **(b)** Locations of selected amino acids across the surface of Sr50’s leucine-rich repeat (LRR) domain. **(c and d)** Predicted structure of AvrSr50 in the identical orientation. **(c)** Positively charged amino acids are indicated in blue with additional residues shown to be important for interactions with Sr50. **(d)** Estimated electrostatic potential is mapped to the surface of AvrSr50. **(e)** Cell death phenotypes on *Nicotiana benthamiana*. Indicated pairs of receptors and effectors were co-infiltrated, and the hypersensitive response (HR) intensity was quantified at three days post infiltration. The statistical significance was accessed with one-way ANOVA followed by a post-hoc Tukey Honestly Significant Difference test. Groups sharing the same letter are not significantly different (*P* > 0.05). Representative HR images corresponding to the median intensity are displayed. **(f)** Simplified schematic representation of the proposed interaction interface between Sr50 and AvrSr50. Amino acids are shown in circles, with their positions labeled adjacent to each residue for Sr50 on green lines and AvrSr50 on red lines. Blue dashed lines indicate proposed interacting pairs. **(g and h)** Predicted poses of Sr50 and AvrSr50 in Model III and IV. Some parts of a loop between β2–β3, mostly including an unstructured region, are removed from AvrSr50 for visualization (positions 42-66). **(i and j)** Superposition of AvrSr50 in Model II and III as well as III and IV. The entire complex structures of the two models were superposed to fix the coordinates of Sr50 consistent. The AvrSr50 structures are then visualized without positions 42 to 66. **(k)** Local environment around Sr50’s K711 in Model IV.

Based on these observations, we postulated that AvrSr50’s R128 would interact with Sr50’s D641 and D643, which are spatially adjacent based on Model II (Fig. 4a). We created AvrSr50^R128E^ and AvrSr50^R129E^ for comparison and co-infiltrated them with Sr50^D641R^ or Sr50^D643K^. Experiments indicated that the mutation at these receptor positions attenuate AvrSr50 recognition (Fig. 4e; Table S1). AvrSr50^R128E^ generated a variable phenotype from no cell death to strong HR in co-infiltration with Sr50. Nevertheless, AvrSr50^R128E^ could enhance HR mediated by Sr50^D641R^ and clearly restore HR with Sr50^D643K^. This suggested that AvrSr50’s R128 interacts with Sr50’s D643 and potentially with D641 (Fig. 4f).

### Refining structural hypotheses with experimental constraints and AlphaFold

To refine our structural hypotheses, we incorporated experimentally identified receptor-effector interactions as restraints. These included D643-R128, K824-E117 and D119, and R904-Q90, E117 and D119 (Fig. 4f). We selected D643 over D641, as these residues were proximal (Fig. 4a), but D643 showed a stronger contribution to immune activation. Compared to Model II, Model III slightly repositioned AvrSr50, improving the fit of its terminal α-helix into the groove of the Sr50 LRR domain and resolving structural abnormality in the previous model (Fig. 4g and 4i).

Although Model III provided a plausible interaction model, we were uncertain whether ColabDock had introduced other structural artifacts, such as backbone clashes observed in Model II. We used ColabFold to eliminate abnormal structural features and remodel the flexible loop regions, using Model III as a structural template. This refined structure, Model IV, closely resembled Model III, with only minor adjustments of AvrSr50’s position (Fig. 4h and 4j). Notably, in Model IV, AvrSr50’s terminal α-helix, enriched with positively charged residues, remained positioned near Sr50’s D643 and K711. However, AvrSr50’s R128 was not directly oriented to interact with Sr50’s D643 (Fig. 4k), suggesting that mutations at these positions may influence effector positioning or that Model IV still retains some structural inaccuracies.

### Multiple Sr50^K711D^ double mutants enhance cell death against AvrSr50^QCMJC^

With increased confidence in our structural hypothesis, we aimed to improve the recognition specificity of Sr50^K711D^. Model IV placed AvrSr50’s Q121 near Sr50’s LRR 13 and 14 (Fig. 5a), identifying W822, E847 and N849 as additional surface-exposed residues to target for engineering towards enhanced effector recognition. These residues were substituted with negatively charged aspartic and glutamic acids, as well as polar asparagine and glutamine, except for N849Q. Among the tested double mutants, Sr50^K711D/W822N^, Sr50^K711D/W822Q^, Sr50^K711D/E847D^, Sr50^K711D/N849D^ and Sr50^K711D/N849E^ could induce HR upon co-expression with AvrSr50^QCMJC^ (Fig. 5b; Fig. S10-11; Table S1). The HR intensity of these double mutants was significantly greater than Sr50^K711D^ (Fig. S11), indicating successful improvement of AvrSr50^QCMJC^ recognition. Moreover, these mutations did not abolish AvrSr50 recognition. Therefore, after several iterations, Model IV allowed successful engineering of Sr50 variants with robust recognition of the AvrSr50^QCMJC^ escape variant.

**Figure 5.**
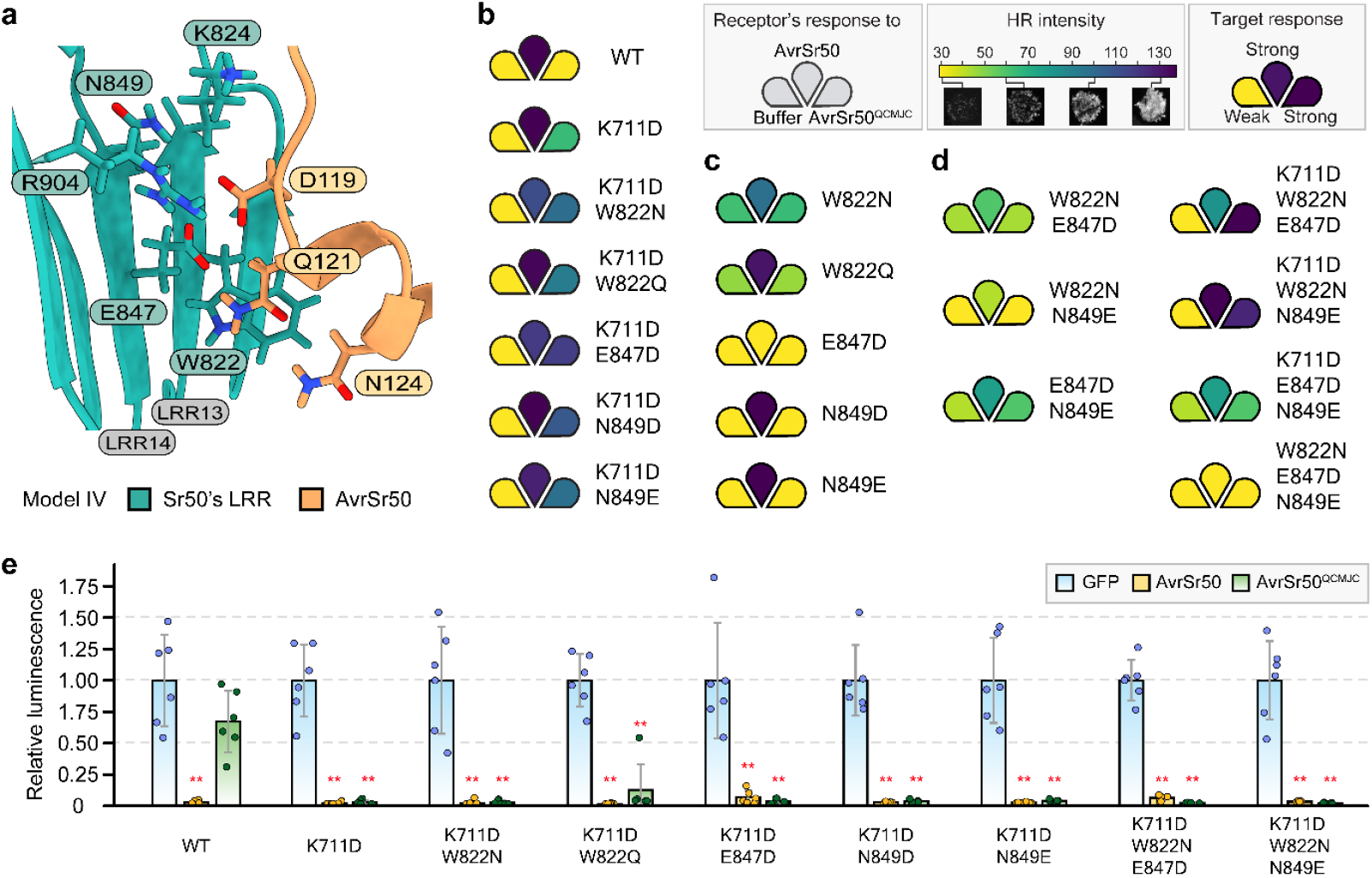
Engineered double and triple Sr50 mutants induce robust HR upon recognition of AvrSr50^QCMJC^. **(a)** Local environment around AvrSr50’s Q121 in Model IV. **(b-d)** Immune responses of receptor variants. Each petal indicates a receptor’s hypersensitive response (HR) to buffer (autoactivity), AvrSr50 and AvrSr50^QCMJC^. HR intensity ranges from 30 to 137, corresponding to the average intensity measured for HR in *Nicotiana benthamiana* co-infiltration with Sr50 and buffer or AvrSr50, respectively. The target response is characterized by no autoactivity and strong HR against both AvrSr50 and AvrSr50^QCMJC^. **(e)** Wheat protoplast cell death assays. Indicated receptors, effectors and luciferase were co-transfected into wheat protoplasts. Luminescence values were normalized using the median of GFP co-transfections for the given receptor. Statistical significance was assessed using the Kruskal-Wallis test followed by pairwise Wilcoxon tests with false discovery rate correction. Significant decreases in luciferase activity compared to GFP controls are denoted (**: *P* < 0.005).

### The K711D mutation is required and synergistic for AvrSr50^QCMJC^ recognition

To further establish relative contributions of the secondary mutations introduced in the Sr50^K711D^ background, we created Sr50^W822N^, Sr50^W822Q^, Sr50^E847D^, Sr50^N849D^ and Sr50^N849E^ single mutants. Sr50^W822N^ and Sr50^W822Q^ led to detectable HR in co-infiltration with AvrSr50^QCMJC^, but the response was associated with an equivalent increase in autoactivity (Fig. 5c; Fig. S10-11). Introducing the K711D substitution back to these two single mutants abolished autoactivity and enhanced HR induced by AvrSr50^QCMJC^ recognition (Fig. 5b). In contrast, the E847D substitution abolished AvrSr50 recognition, while Sr50^N849D^ and Sr50^N849E^ did not alter recognition specificity (Fig. 5c). Adding the K711D substitution to these single mutants also enabled AvrSr50^QCMJC^ recognition while restoring AvrSr50 recognition (Fig. 5b). Therefore, we conclude that while secondary mutations enhance the recognition of Sr50^K711D^, the K711D mutation is essential for AvrSr50^QCMJC^ recognition.

The phenotypic epistasis between the single and double mutants was unexpected, as we initially postulated that the effects of distantly located mutations would be additive. To investigate the interplay between the targeted residues, we generated all combinations of double and triple mutants involving K711D, W822N, E847D and N849E (Fig. 5c-d; Fig. S10-11). Among the four triple mutants, Sr50^K711D/W822N/E847D^ and Sr50^K711D/W822N/N849E^ induced stronger HR when co-expressed with AvrSr50^QCMJC^ than the double mutants (Fig. S11). Interestingly, Sr50^W822N/E847D^ and Sr50^W822N/N849E^, without the K711D substitution, showed impaired receptor activation by AvrSr50 and did not recognize AvrSr50^QCMJC^ (Fig. 5d). Collectively, our results indicated that the K711D substitution underlies robust AvrSr50^QCMJC^ recognition and functions synergistically with other mutations.

### Engineered elite Sr50 receptor variants mediate AvrSr50^QCMJC^ recognition in transient expression assays in wheat

To evaluate the efficacy of engineered elite Sr50 mutants against AvrSr50^QCMJC^ in wheat, we conducted wheat protoplast-based cell death assays. Effector-mediated Sr50 activation was assessed by cell death analysis using luciferase activity as proxy for cell viability. Luminescence upon co-transfection of each Sr50 variant with AvrSr50 and AvrSr50^QCMJC^ was compared to that observed with GFP as a control. Expression of all tested receptors together with AvrSr50 led to robust HR (Fig. 5e). While Sr50 could not induce cell death when co-expressed with AvrSr50^QCMJC^, all examined receptor mutants successfully triggered HR (Fig. 5e). The magnitudes of immune responses were similar across all mutants, which we attribute to the use of the strong constitutive *Zea mays* ubiquitin promoter (pZmUBQ), as observed for p35S in the respective *N. benthamiana* assays (Fig. 1).

### Engineered Sr50 alleles overlap with highly variable amino acid sites essential for effector binding

LRR residues with high sequence variation among homologous sequences were shown to be associated with effector recognition (Prigozhin and Krasileva, 2021). To quantify natural diversity at LRRs, we calculated normalized Shannon entropy for the Sr50 homologous group curated from the previous study (Tamborski *et al*., 2023). Our analysis confirmed that the central effector binding site identified in our computational structure-guided experiments without knowledge of the natural diversity data overlaps with highly variable LRR (hvLRR) residues. Notably, a patch of hvLRRs with the highest Shannon entropy was located at the terminal LRRs above the central beta sheets (Fig. 6a). This region includes K824, W822N, E847, N849 and R904, all critical for effector recognition. Highly variable K711 was positioned more closely to non-hv D641 and D643 (Fig. 6a). Small variations in these two positions may be attributed to their proximity to the NB-ARC domain, where incompatible mutations could disrupt interdomain interactions.

**Figure 6.**
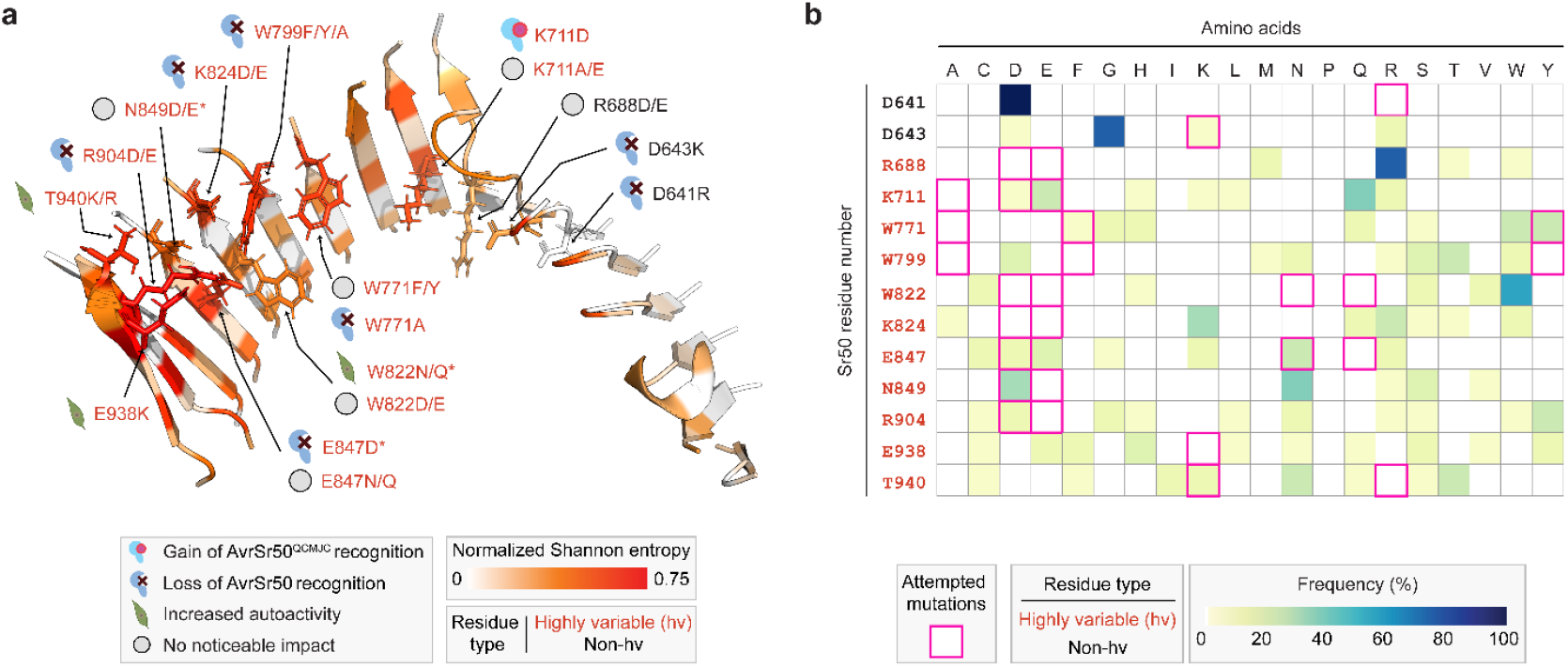
Structural and evolutionary features of mutagenized receptor residues. **(a)** Inner beta strands of leucine-rich repeat (LRR) domain of Sr50. The residues are colored based on normalized Shannon entropy scores, which indicate variability of homologous sequences within the Sr50 family. The score ranges from 0 (no sequence variability) to 1 (complete variability). The score was capped at 0.75 for visualization. Scores greater than 0.347 indicate highly variable (hv). The positions chosen for mutagenesis and the phenotypes of single mutants are indicated. The mutations indicated with asterisks lead to different phenotypes when K711D substitution is introduced together. **(b)** Frequency of amino acids within the Sr50 family in the given homologous positions. The mutations introduced in this study are indicated as pink boxes.

Adjacent to the hvLRR patch, two tryptophan residues, W771 and W799, were initially hypothesized to be important for affinity towards AvrSr50 (Fig. 1e and 6a). Among these, W799, located the closest to the hvLRR patch, was found to be essential for AvrSr50 recognition and could not be replaced by tyrosine or phenylalanine (Fig. S12). In contrast, W771 could tolerate these substitutions (Fig. S12). Interestingly, in AvrSr50^Q121K/N124W^, a mutant that enhanced immune responses for Sr50^K711D^ (Fig. 2), the introduced tryptophan lies near W771, suggesting that the increased HR likely resulted from increased receptor-effector affinity (Fig. S13). Overall, our structural model underscores the importance of the terminal hvLRR patch for interaction with AvrSr50’s central binding site, particularly involving residues D119 and Q121, the mutations of which were shown to induce recognition escape (Fig. 2 and 3). These findings highlight the rapid co-evolution at the receptor-effector interaction interface.

### Engineered Sr50 variants extend beyond natural diversity

Specific amino acid substitutions that we introduced to create functional double and triple receptor mutants were rare among the Sr50 homologs (Fig. 6b). For instance, aspartic acid appeared in 6.25% of instances at position 711, while asparagine and glutamine were absent at position 822. Similarly, aspartic acid appeared in 12.5% at position 847, and glutamic acid in no instances at position 849. Aspartic acid was relatively frequent at position 849, appearing in 30% of sequences. Notably, no homologous sequences contained the specific combinations of amino acids required for AvrSr50^QCMJC^ recognition. Collectively, while our structure-guided engineering targeted regions of high natural diversity, the resulting Sr50 alleles were distinct from known natural variations, underscoring their novelty. Ability to successfully engineer new functional combinations of amino acids that are distinct from allelic variation present in natural diversity has potential implications in receptor durability. Since our engineered Sr50 variants might not yet exist in nature, the pathogen has not been subjected to selective pressure to overcome them.

### The K711D substitution alters AlphaFold 2 predictions

Our initial attempt to model the Sr50–AvrSr50 complex with AF2-Multimer resulted in low quality predictions that hindered confident engineering of Sr50 (Fig. 7a). We tried modeling the complex structure of Sr50^K711D/W822N/E847D^ and AvrSr50^QCMJC^, as the engineered receptor mutant induced strong HR when co-expressed with the effector. Unexpectedly, AF2 generated a model with a confidence (ipTM) score of 0.609 that closely matched Model IV (Fig. 7a; Table S2). Similarly, a high confidence model was produced for the Sr50^K711D/W822N/E847D^ and AvrSr50 complex with an ipTM score of 0.798.

**Figure 7.**
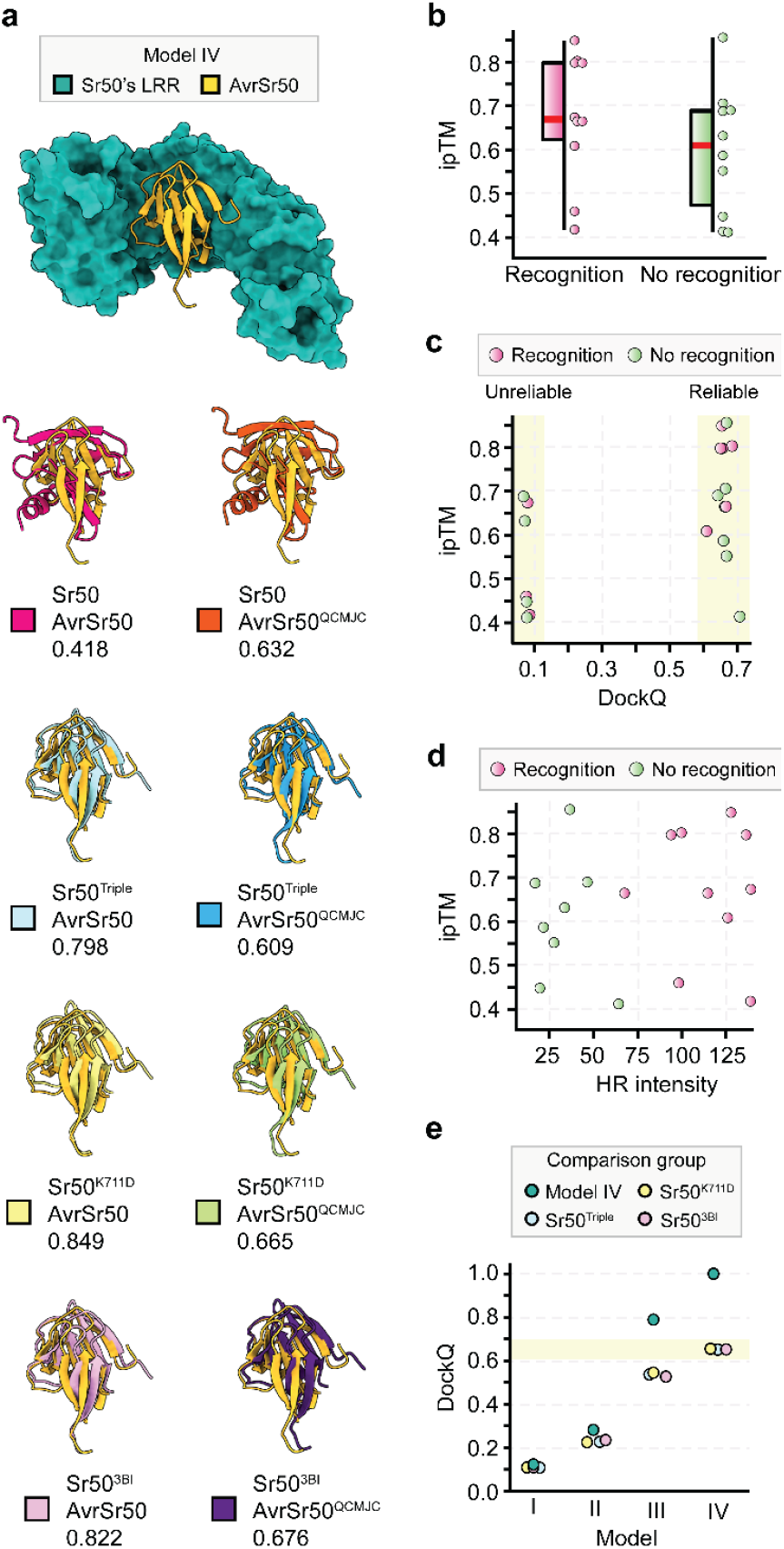
AlphaFold confidence scores distinguish reliable and unreliable predictions but do not fully correlate with immune activation outcomes phenotypes. **(a)** Structural superposition of AvrSr50 in AlphaFold 2 (AF2) models compared to Model IV. Predicted protein complex structures were superposed against Model IV to keep receptor coordinates consistent. The AvrSr50 structures were then visualized. Labels indicate the receptors and effectors used for prediction and their associated confidence scores (ipTM). Sr50^Triple^ refers to Sr50^K711D/W822N/N849E^. **(b)** Distribution of ipTM confidence scores based on immune outcomes. **(c)** Distribution of ipTM confidence scores in relation to DockQ scores. Each prediction model was compared to Model IV, and a DockQ score was computed to assess prediction accuracy. DockQ above 0.6 indicates that models are reliable. **(d)** Distribution of ipTM confidence scores in relation to hypersensitive response (HR) intensity. **(e)** Comparison of four structural hypotheses (Models I to IV) to the final structural hypothesis (Model IV) and AF2 models. Sr50^Triple^ refers to Sr50^K711D/W822N/N849E^. DockQ scores can be originally divided into incorrect (0 < DockQ < 0.23), acceptable quality (0.23 ≤ DockQ < 0.49), medium quality (0.49 ≤ DockQ < 0.80) and high quality (0.80 ≤ DockQ ≤ 1). In comparison between Model IV and AF2 models, the highest similarity resulted in the DockQ scores between 0.6 and 0.7, which were classified as reliable models.

To identify the specific sequence variation responsible for altering AF2’s predictive behavior, we modeled the complex structures of Sr50^K711D^, Sr50^W822N^ and Sr50^E847D^ with AvrSr50. Only the K711D substitution, but not W822N and E847 substitutions, altered AF2’s behavior (Fig. 7a and S14), resulting in a high confidence model with a score of 0.849. Collectively, these results indicated that the single K711D substitution not only modified Sr50’s biological response to the effector but was also sufficient to influence AF2’s structural prediction outcome.

A recent study modeled the structure of Sr50^3BI^ and AvrSr50 with AF2 to predict the effector binding site (Gómez De La Cruz *et al*., 2024). Sr50^3BI^ differs from Sr50 by 12 amino acids with a short stretch of amino acids transferred from the barley NLR MLA3 but does not contain mutations at position 711. We found that the Sr50^3BI^ and AvrSr50 structure showed good agreement with Model IV as well as the AF models of Sr50^K711D^ and Sr50^K711D/W822N/E847D^ complexes (Fig.7a). These results indicated that slight changes in the input sequences can alter the behavior and outcome of AF2 prediction. This potentially suggests that iterative *in silico* mutagenesis of input sequences could aid reliable prediction of NLR-effector complexes.

### AlphaFold confidence scores separate reliable and unreliable predictions but do not fully correlate with immune activation outcomes

To assess the applicability of AF-guided predictions towards future Sr50 engineering, we examined the relationship between AF2-Multimer’s ipTM confidence scores and immune phenotypes by analyzing selected receptor-effector mutant pairs from our study (Table S2). An ipTM score above 0.8 is generally associated with high-confidence prediction of protein complexes, while models with scores between 0.6 and 0.8 could be either correct or incorrect (Evans *et al*., 2021; Wee and Wei, 2024). We observed that while receptor-effector pairs that elicited immune responses often had higher ipTM scores, this trend was not statistically significant (Fig. 7b). To further evaluate the predictive power of AF2, we compared each AF2 model to Model IV, assessing its accuracy as DockQ scores. We used DockQ > 0.6 as the threshold for reliable prediction. As previously suggested (Wee and Wei, 2024), all models with ipTM scores above 0.8 were reliable, whereas models with ipTM scores between 0.6 and 0.7 varied in accuracy, yielding both reliable and unreliable predictions (Fig. 7c). However, a receptor-effector pair predicted with an ipTM score of 0.856 did not elicit immune responses (Table S2), demonstrating that even high-confidence structural predictions do not always translate to functional interactions. Similarly, we did not find any clear correlation between confidence scores and the magnitude of immune responses (Fig. 7d). These findings highlight both the utility and limitations of AF-guided NLR engineering. While the ipTM scores can distinguish reliable predictions from unreliable ones, structural predictions alone are insufficient for determining immune activation, as incompatibility at the residue level remains a critical factor.

### All models are wrong, but some are useful

Our workflow for predicting Sr50–AvrSr50 interactions demonstrated that, at present, modeling-based engineering approaches need to be iterative. Through three rounds of modeling and a round of computational refinement, we arrived at our final solution, Model IV. While the true accuracy of the model can only be evaluated when Cryo-EM structure becomes available, Model IV closely resembled independently derived AF models (Fig. 7a). Comparing all modeling iterations to Model IV and other AF models, we observed that our initial structural hypothesis (Model I) was completely incorrect (Fig. 7d). However, in each refinement cycle, experimental constraints and structural analyses progressively improved the accuracy of our models. This iterative process highlights that while no model is perfect, each step contributes valuable insights toward the next.

## Discussion

Experimentally determined NLR-effector complex structures offer NLR engineering strategies; however, their uses currently remain limited. Predicting the effects of mutations in pathogen effector genes on NLR-mediated recognition and host disease phenotypes has been a major challenge hindering engineering efforts. Therefore, accumulating evolutionary and experimental data is crucial for effective engineering solutions. Our data demonstrates that iterative computational modeling together with experimental determination of structural constraints is an effective strategy for inferring NLR-effector structures and restoring recognition of the escape effector mutants. Although our initial structure hypothesis contained inaccuracies, iterative structural refinements progressively improved the predicted models, eventually enabling the engineering of elite Sr50 variants against naturally occurring AvrSr50^QCMJC^. As such, this study provides a path to expand the recognition specificities of NLR genes to previously unrecognized pathogen effector variants. Deployment of engineered receptors in elite wheat cultivars holds exciting potential to broaden Sr50-mediated resistance specificity to *Pgt* isolate QCMJC and other isolates carrying AvrSr50^QCMJC^ that have evaded Sr50 recognition (Chen *et al*., 2017; Mago *et al*., 2015).

Our approach differs from two recently showcased LRR engineering approaches (Gómez De La Cruz *et al*., 2024; Lawson *et al*., 2024). Lawson et al. (2024) expanded recognition specificity of the barley NLR MLA7 to recognize AVR_A13_. They leveraged the experimentally determined structure of MLA13–AVR_A13_-1 complex, combined with sequence comparison of the two closely related functional receptors MLA7 and MLA13. This structural and sequence information allowed transferring a single key residue involved in effector binding to MLA7. Similarly, Gómez De La Cruz et al. (2024) engineered Sr50 to recognize an additional effector, Pwl2, from *Maganaporthe oryzae*, which is known to target HIPP43 in rice. The identification of the Pwl2 binding site in the NLR MLA3 was guided by AF2, yet the existing crystal structure of the HIPP43–Pwl2 complex confirmed the similarity between the two binding sites (Brabham *et al*., 2024; Zdrzałek *et al*., 2024). Their approach focused on transferring this interface to Sr50 while retaining its ability to recognize AvrSr50. In contrast, we engineered Sr50 variants to recognize an effector undetectable by any known NLRs, distinguishing our approach from these previous studies. While our work greatly benefited from the crystal structure of AvrSr50^QCMJC^ and the identification of its Q121K escape mutation (Ortiz *et al*., 2022), our strategy was developed without prior knowledge of any natural receptors recognizing AvrSr50^QCMJC^. Remarkably, our engineered Sr50 variants, despite their robust ability to induce HR in response to AvrSr50^QCMJC^ in both *N. benthamiana* and wheat protoplast, are absent among sequenced genomes, highlighting their novelty.

A critical finding of our study is how epistasis among amino acids at the LRR ligand binding site influences phenotypic outcomes and evolutionary trajectories (Fig. 8). Epistasis arises when a mutation alters the effect of other mutations, thereby reshaping accessible evolutionary pathways and evolutionary outcomes (Starr and Thornton, 2016). In the interaction with *Pgt*, wheat may first accumulate substitutions like E847D or N849E, which leads to loss of function or does not alter recognition, respectively (Fig. 8a; I and II). These mutations would likely be selected against, due to the lack of AvrSr50^QCMJC^ recognition, and subsequent acquisition of beneficial mutations like K711D may not occur. In contrast, mutations conferring autoactivity at the cost of development (e.g. Sr50^W822N^) or weak-to-intermediate recognition (e.g. Sr50^K711D^) could facilitate further modifications that either enhance HR or reduce autoactivity (Fig. 8a; III and IV). Among tested mutations, K711D as the initial mutation offers the greatest potential for acquiring robust resistance, as its emergence modifies the functional effects of subsequent mutations. Our results highlight that due to epistasis, the order in which mutations arise can determine evolutionary trajectories, either restricting or expanding adaptive potential, even when different mutational paths ultimately converge on the same outcomes (Fig. 8b).

**Figure 8.**
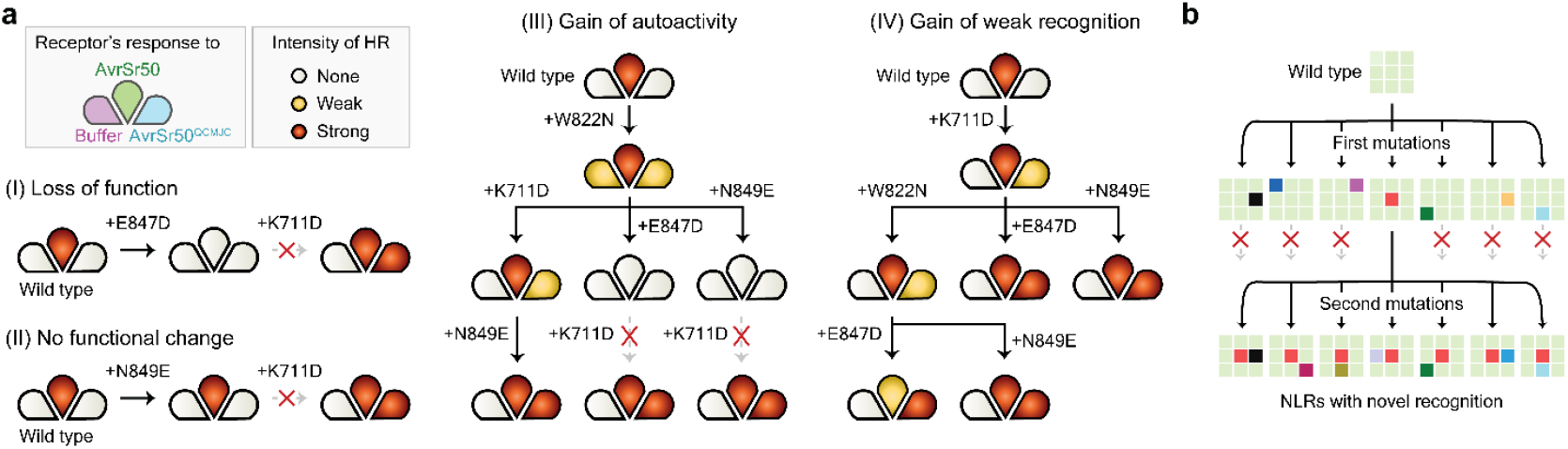
The initial mutation influences subsequent evolutionary opportunities. **(a)** Phenotypes of engineered Sr50 mutants along potential mutational paths. Each petal indicates a receptor’s hypersensitive response (HR) to buffer (autoactivity), AvrSr50 and AvrSr50^QCMJC^. Starting from the wild type, Sr50 accumulates a single mutation along each path. The path eventually leads to reduced autoactivity or enhanced HR against AvrSr50^QCMJC^. Dotted lines indicate paths that become unavailable due to specific phenotypic outcomes of introduced mutations. **(b)** The influence of the initial mutations in subsequent mutational opportunities. A colored box indicates a unique mutation. The first mutation can either limit or expand subsequent mutational trajectories, despite leading to an identical set of mutations.

Our findings also advance our understanding of AlphaFold’s capabilities and limitations. While AF cannot model the complex structure of Sr50 and AvrSr50, AF2 produced high-confidence models of Sr50 variants and AvrSr50 (Fig 7). Notably, the single K711 substitution rectified the previously unsuccessful prediction, altering both biological and computational outcomes. While AF2 is not specifically trained to predict the impact of amino acid mutations in protein folding, our results suggest that even minor changes in input sequences can significantly influence the outcome of protein complex structure prediction. Furthermore, our data, derived from 6,000 infiltration quantifications, provides a robust representation of diverse biological responses between Sr50 and AvrSr50 variants (Table S1). This resource will inform further correlations between AF’s prediction confidence, model reliability and immune outcomes, also serving as an experimentally derived training dataset for emerging computational approaches. These insights will open opportunities to ‘hack’ AF2 for *in silico* screening of receptor-effector.

We postulate that K711 likely plays a supplementary role in Sr50’s interactions with AvrSr50, as mutations at this position did not significantly affect AvrSr50 recognition (Fig. 1 and S5). However, when AvrSr50^QCMJC^ disrupts the interaction with Sr50’s hvLRRs through its K121 (Fig. 3 and 6), the K711D substitution would facilitate additional interactions with AvrSr50^QCMJC^’s alpha helix enriched with positively charged residues (Fig. 4). While the HR induced by Sr50^K711D^ in response to AvrSr50^QCMJC^ was only weak to intermediate in *N. benthamiana* (Fig. 1 and 2), this subtle interaction likely enables proper alignment of AvrSr50^QCMJC^ to Sr50, allowing additional receptor mutations to further stabilize the interaction (Fig. 5). This mechanism may explain why the K711D substitution is required for AvrSr50^QCMJC^ recognition but also synergistic with additional mutations. Given the challenges of protein-level engineering due to epistasis and complexity of protein-protein interactions, expression engineering approaches offer an appealing alternative. Weak recognition can be developed first through point mutations, and fine-tuned expressions can minimize autoactivity while maximizing pathogen-induced responses (Fig. 1). In nature, NLR expression evolution might complement protein evolution to gain new resistance, as extensive intraspecies diversity of NLR expression was observed (Prigozhin *et al*., 2024).

Expanding recognition specificity of Sr50 to other paralogs and sequence-unrelated structurally similar (SUSS) effectors is the next challenge (Seong and Krasileva, 2021, 2023). While AvrSr50 represents a SUSS effector family, no other members are yet known to interact with NLRs. Alternatively, MLA receptors from barley and their effectors from *Blumeria* species offer compelling systems (I. M. Saur *et al*., 2019). Not only do MLAs share close evolutionary relationships with Sr50 (Tamborski *et al*., 2023), but also they recognize sequence-divergent or SUSS effectors from an expanded RNAse-like protein family with rich structural data (Cao *et al*., 2023; Lawson *et al*., 2024; Pennington *et al*., 2019; Seong and Krasileva, 2023). Leveraging existing data can aid in hypothesizing initial complex structures and predicting interactions. Elucidating the structures of Sr and MLA receptors with their effectors can reveal how NLRs have evolved to detect SUSS effectors, and the framework of this study may guide experimental designs and engineering solutions for NLR-SUSS effector systems.

## Methods

### Vectors and mutagenesis

We used the previously generated constructs (Tamborski *et al*., 2023). The receptor constructs carried *Sr50* under the *pRPP13* promoter and the *tNos* terminator, or the *p35S* promoter and the *t35S* terminator. The effector constructs contained *AvrSr50* and *AvrSr50*^*QCMJC*^ with the *p35S* promoter and *t35S* terminator. All additional mutations were introduced with QuikChange Lightning Site-directed Mutagenesis Kits from Agilent. We followed the standard protocol but reduced the volume of all reagents by four. All primers and mutants used in this study are available in Table S3.

### Transformation and mutation confirmation

XL10-Gold ultracompetent cells were transformed, following the standard protocol given in QuikChange Lightning Site-directed Mutagenesis Kits. Plasmids were extracted from a liquid culture inoculated with a single colony, following the standard protocol of The QIAprep Spin Miniprep Kit. The desired mutation was confirmed with Sanger sequencing.

About 40 μL of *Agrobacterium tumefaciens* GV3101:pMP90 was mixed with 100 ng of the mutagenized plasmid in a 1.5 ml plastic tube and transferred to an electroporation cuvette. After an electric pulse, the cells were transferred back to the tube with 250 μL of LB media and shaken at 28°C and 250 rpm for two hours. The liquid culture was plated on LB agar containing Carbenicillin, Rifampicin and Gentamicin, and the plates were incubated at 28°C for two days. A single colony was selected and transferred to a liquid LB medium and grown for a day.

We additionally confirmed the mutations in the transformed *Agrobacterium*. Each mutant was re-plated from its glycerol stock on LB agar containing Carbenicillin, Rifampicin and Gentamicin and grown at 28°C for two days. Colonies were scraped from the plates, resuspended in 10 μL of water and incubated at 98°C for 10 minutes. The cells were centrifuged for one minute, and target regions with introduced mutations were amplified, following the PCR Protocol for Phusion® High-Fidelity DNA Polymerase (M0530) or for repliQa HiFi ToughMix. The mutation was confirmed with Sanger sequencing.

### *Agrobacterium*-mediated transient gene expression in *N. benthamiana*

The liquid cultures containing *Agrobacterium* transformants were centrifuged at 6,400 g for five minutes, and the pellets were resuspended in infiltration medium composed of deionized water, 10 mM MES (pH 5.6), 10 mM MgCl_2_, 150 μM acetosyringone. The optical density (OD_600_) of each transformant was re-adjusted to 0.6. The transformants carrying receptors and effectors were mixed to adjust the OD_600_ of the receptors to 0.1 (for p35S) or 0.3 (for *pRPP13*). The OD_600_ of the effectors was set to 0.3. Leaves of 4-5 weeks old *N. benthamiana* were inoculated with the suspension using blunt syringes. The phenotypes were recorded at 2 days post infiltration (dpi) for the *p35S* promoter and at 3 dpi for the *pRPP13* promoter.

### Cell death quantification and statistics

The infiltrated *N. benthamiana* leaves were imaged with the ChemiDoc MP Imaging System and Image Lab v5.2.1 (https://www.bio-rad.com). Following the previous publication (Landeo Villanueva *et al*., 2021), we used green epiillumination with a filter set to 605/50. Exposure time was set to 0.5 seconds. To quantify the cell death, we manually selected the treated area and measured the mode of intensity with ImageJ v2.14.0 (Rueden *et al*., 2017). The statistics were computed in R v4.1.3 (Ihaka and Gentleman, 1996).

### Cell death assays in wheat protoplasts

In wheat protoplast assays, all gene expression were driven by pZmUBQ and the nopaline synthase terminator (tNos) with the pICH41421 vector backbone (Tamborski *et al*., 2023). Experiments were performed according to the previous study (I. M. L. Saur *et al*., 2019) with the modification that transfection volumes for plasmid DNA were 8 μL (4 μg) for *pZmUBQ:luciferase*, 10 μL (10 μg) for *Sr50* variants, and 10 μL (10 μg) for *GFP* as well as *AvrSr50* variants.

### Protein extraction and immunoblotting assays

Six *N. benthamiana* leaf discs of 0.8 cm in diameter were collected at 1 dpi. The samples were frozen in liquid nitrogen and ground with a bead beater at 1,500 Hz for 1 min with two 3.2 mm stainless beads. Protein was extracted with 300 μL of the 2x Laemmli sample buffer (Biorad) with 5% β-mercaptoethanol. Samples were boiled at 95°C for 5 min and centrifuged at maximum speed for 10 min at 4 °C. Supernatants were transferred into fresh tubes for SDS PAGE analysis. 10 μL of protein extraction was separated on 15% Mini-PROTEAN® TGX™ Precast Protein Gels (Biorad, 15-well), transferred to PVDF membrane (Biorad) at 300 mA for 70 min. Immunoblotting was performed using rat HRP-conjugated α-HA (monoclonal 3F10, Roche) and subsequently chemiluminescent substrate SuperSignal™ West Pico PLUS (Thermo Scientific™). Total protein was stained using Ponceau S.

### Protein structure prediction and visualization

The initial Sr50 and AvrSr50 structures were predicted by AlphaFold v2.2.2(Jumper *et al*., 2021), with the full database, available homologous templates and model_preset set to monomer. Protein complexes were predicted with ColabFold v1.5.2 that relies on AlphaFold v2.3.1, with alphafold2_multimer_v3 and template_model set to pdb100, as well as AlphaFold 3 (Abramson *et al*., 2024; Evans *et al*., 2021; Mirdita *et al*., 2022). We used customized ColabDock through Google Colab to obtain Models II and III (Feng *et al*., 2023). Pairwise constraints derived from the experiments were provided. Among the five models, the best structure (1st_best) was used for the analysis. The side chains of this structure were relaxed with Amber, a module within ColabFold.

The AF2 models, which were compared to Model IV, were generated with ColabFold v1.5.5. The structure of AvrSr50^QCMJC^ was submitted as a template (PDB:7MQQ) (Ortiz *et al*., 2022). Five models were predicted with the alphafold2_multimer_v3 model, and the best structure was relaxed. The default parameters were used, except for num_recycles set to 24 and pair_mode changed to unpaired. We used PyMOL v2.5.2 and ChimeraX v1.9 for protein structure superposition and visualization (Jurrus *et al*., 2018; Pettersen *et al*., 2021; The PyMOL Molecular Graphics System). Structural similarities between complex structures were quantified with DockQ v2.1.1(Basu and Wallner, 2016).

### Molecular docking and initial model selection

The best Sr50 and AvrSr50 monomer models were submitted to ZDOCK, HDOCK and ClusPro web servers as a receptor and a ligand, respectively (Kozakov *et al*., 2017; Pierce *et al*., 2014; Yan *et al*., 2020). From each server, 100 models were obtained, and each model was evaluated for the following criteria, based on the backbone distances (C_β_ or C_α_ of glycine). First, all Sr50 residues in the coiled-coil and NB-ARC domains (positions 1 to 520) are not within 8 Å of AvrSr50. Second, AvrSr50 should touch the NB-ARC latch–a loop structure exposed from the NB-ARC domain, forming close contact with LRRs (positions 492-499). Specifically, Sr50’s E494 is within 12 Å of AvrSr50. This residue was chosen as the predicted Sr50 structure suggested that its sidechain points toward the putative effector binding site surrounded by the concave of LRR units. The distance cut-off was relaxed based on our assumption that the interaction between the NB-ARC latch and the effectors occur through long side chains. Third, AvrSr50’s Q121 is within 8 Å of Sr50’s LRR domain (Ortiz *et al*., 2022; Tamborski *et al*., 2023). Fourth, at least 12 hvLRR residues on the inner β-strands or upper loops of Sr50 are within 8 Å of AvrSr50 (Prigozhin and Krasileva, 2021). Then, from the models that satisfied these criteria, the effector poses were clustered with the RMSD (root mean square deviation) cut-off of 3.0. We required at least two predictors to produce similar poses, despite their differing scoring functions and underlying algorithms. A representative conformation was chosen from each cluster based on the model ranking.

### Evolutionary analyses

The multiple sequence alignment of MLA family members and the normalized Shannon entropy for the Sr50 homologous group were obtained from the previous study and analyzed with the identical workflow (Tamborski *et al*., 2023).

## Data availability

All scripts and command lines used for computational analyses are available and recorded in Github: https://github.com/s-kyungyong/Sr50_AvrSr50/. All input, intermediate and output data were deposited in Zenodo: https://zenodo.org/doi/10.5281/zenodo.13205869

## Acknowledgements

We thank China Lunde for phenotyping, and Sarah Weber for attempting transient gene expression assays. We thank Chandler Sutherland, China Lunde and Dr. Daniil Prigozhin for the critical review of the manuscript. K.S. is supported by the Berkeley BioEnginuity Fellowship. I.ML.S. and S.C.S. have been supported by the German Research Foundation (DFG) Emmy Noether Programme (SA 4093/1-1 to I.M.L.S.) and the DFG Collaborative Research Centre Grant (SFB-1403 – Project-ID 414786233 to I.M.L.S.). K.V.K. has been supported by the Gordon and Betty Moore Foundation (grant number: 8802) as well as the joint funding from the Foundation for Food and Agriculture and 2Blades (CA19-SS-0000000046), the Innovative Genomics Institute and the National Institute of Health Director’s Award (1DP2AT011967-01).

## Contributions

K.S. conceptualized and designed the project. K.S. wrote the manuscript with edits from K.V.K. and W.W. W.W. performed Western blot and cloning. S.C.S. and I.ML.S. conducted protoplast assays. K.S. performed computational analyses. K.S., W.W., B.V., A.D., G.R., R.K. and L.P. conducted all other experiments. K.V.K. supervised the research.

## Competing interests

The authors declare no competing interests.

## Notes

### Competing Interest Statement

The authors have declared no competing interest.

### Summary of Updates

This revision contains wheat protoplast data, which confirms that engineered Sr50 variants are also functional in the transient gene expression assays in wheat. Figures and texts were improved for clarity.

## References

Abramson, J., Adler, J., Dunger, J., Evans, R., Green, T., Pritzel, A., et al. (2024) Accurate structure prediction of biomolecular interactions with AlphaFold 3. Nature.

Arora, S., Steuernagel, B., Gaurav, K., Chandramohan, S., Long, Y., Matny, O., et al. (2019) Resistance gene cloning from a wild crop relative by sequence capture and association genetics. Nat Biotechnol, 37, 139– 143.

Baggs, E., Dagdas, G., and Krasileva, K. (2017) NLR diversity, helpers and integrated domains: making sense of the NLR IDentity. Current Opinion in Plant Biology, 38, 59–67.

Basu, S. and Wallner, B. (2016) DockQ: A Quality Measure for Protein-Protein Docking Models. PLoS ONE, 11, e0161879.

Bentham, A.R., De La Concepcion, J.C., Benjumea, J.V., Kourelis, J., Jones, S., Mendel, M., et al. (2023) Allelic compatibility in plant immune receptors facilitates engineering of new effector recognition specificities. The Plant Cell, 35, 3809–3827.

Brabham, H.J., Gómez De La Cruz, D., Were, V., Shimizu, M., Saitoh, H., Hernández-Pinzón, I., et al. (2024) Barley MLA3 recognizes the host-specificity effector Pwl2 from Magnaporthe oryzae. The Plant Cell, 36, 447–470.

Cao, Y., Kümmel, F., Logemann, E., Gebauer, J.M., Lawson, A.W., Yu, D., et al. (2023) Structural polymorphisms within a common powdery mildew effector scaffold as a driver of coevolution with cereal immune receptors. Proc. Natl. Acad. Sci. U.S.A., 120, e2307604120.

Cesari, S., Xi, Y., Declerck, N., Chalvon, V., Mammri, L., Pugnière, M., et al. (2022) New recognition specificity in a plant immune receptor by molecular engineering of its integrated domain. Nat Commun, 13, 1524.

Chen, J., Upadhyaya, N.M., Ortiz, D., Sperschneider, J., Li, F., Bouton, C., et al. (2017) Loss of AvrSr50 by somatic exchange in stem rust leads to virulence for Sr50 resistance in wheat. Science, 358, 1607–1610.

Dangl, J.L., Horvath, D.M., and Staskawicz, B.J. (2013) Pivoting the Plant Immune System from Dissection to Deployment. Science, 341, 746–751.

De La Concepcion, J.C., Franceschetti, M., MacLean, D., Terauchi, R., Kamoun, S., and Banfield, M.J. (2019) Protein engineering expands the effector recognition profile of a rice NLR immune receptor. eLife, 8, e47713.

Dodds, P.N. and Rathjen, J.P. (2010) Plant immunity: towards an integrated view of plant–pathogen interactions. Nat Rev Genet, 11, 539–548.

Evans, R., O’Neill, M., Pritzel, A., Antropova, N., Senior, A., Green, T., et al. (2021) Protein complex prediction with AlphaFold-Multimer. Bioinformatics.

Farnham, G. and Baulcombe, D.C. (2006) Artificial evolution extends the spectrum of viruses that are targeted by a disease-resistance gene from potato. Proc. Natl. Acad. Sci. U.S.A., 103, 18828–18833.

Feng, S., Chen, Z., Zhang, C., Xie, Y., Ovchinnikov, S., Gao, Y., and Liu, S. (2023) ColabDock: inverting AlphaFold structure prediction model for protein-protein docking with experimental restraints. Bioinformatics.

Förderer, A., Li, E., Lawson, A.W., Deng, Y., Sun, Y., Logemann, E., et al. (2022) A wheat resistosome defines common principles of immune receptor channels. Nature, 610, 532–539.

Gómez De La Cruz, D., Zdrzałek, R., Banfield, M.J., Talbot, N.J., and Moscou, M.J. (2024) Molecular mimicry of a pathogen virulence target by a plant immune receptor.

Harris, C.J., Slootweg, E.J., Goverse, A., and Baulcombe, D.C. (2013) Stepwise artificial evolution of a plant disease resistance gene. Proc. Natl. Acad. Sci. U.S.A., 110, 21189–21194.

Huang, H., Huang, S., Li, J., Wang, H., Zhao, Y., Feng, M., et al. (2021) Stepwise artificial evolution of an Sw-5b immune receptor extends its resistance spectrum against resistance-breaking isolates of Tomato spotted wilt virus. Plant Biotechnology Journal, 19, 2164–2176.

Ihaka, R. and Gentleman, R. (1996) R: A Language for Data Analysis and Graphics. Journal of Computational and Graphical Statistics, 5, 299–314.

Jones, J.D.G. and Dangl, J.L. (2006) The plant immune system. Nature, 444, 323–329.

Jumper, J., Evans, R., Pritzel, A., Green, T., Figurnov, M., Ronneberger, O., et al. (2021) Highly accurate protein structure prediction with AlphaFold. Nature, 596, 583–589.

Jurrus, E., Engel, D., Star, K., Monson, K., Brandi, J., Felberg, L.E., et al. (2018) Improvements to the APBS biomolecular solvation software suite. Protein Science, 27, 112–128.

Kozakov, D., Hall, D.R., Xia, B., Porter, K.A., Padhorny, D., Yueh, C., et al. (2017) The ClusPro web server for protein–protein docking. Nat Protoc, 12, 255–278.

Kroj, T., Chanclud, E., Michel-Romiti, C., Grand, X., and Morel, J. (2016) Integration of decoy domains derived from protein targets of pathogen effectors into plant immune receptors is widespread. New Phytologist, 210, 618–626.

Landeo Villanueva, S., Malvestiti, M.C., Van Ieperen, W., Joosten, M.H.A.J., and Van Kan, J.A.L. (2021) Red light imaging for programmed cell death visualization and quantification in plant–pathogen interactions. Molecular Plant Pathology, 22, 361–372.

Lawson, A.W., Flores-Ibarra, A., Cao, Y., An, C., Neumann, U., Gunkel, M., et al. (2024) The barley MLA13-AVR _A13_ heterodimer reveals principles for immunoreceptor recognition of RNase-like powdery mildew effectors.

Ma, S., Lapin, D., Liu, L., Sun, Y., Song, W., Zhang, X., et al. (2020) Direct pathogen-induced assembly of an NLR immune receptor complex to form a holoenzyme. Science, 370, eabe3069.

Mago, R., Zhang, P., Vautrin, S., Šimková, H., Bansal, U., Luo, M.-C., et al. (2015) The wheat Sr50 gene reveals rich diversity at a cereal disease resistance locus. Nature Plants, 1, 15186.

Maidment, J.H., Shimizu, M., Bentham, A.R., Vera, S., Franceschetti, M., Longya, A., et al. (2023) Effector target-guided engineering of an integrated domain expands the disease resistance profile of a rice NLR immune receptor. eLife, 12, e81123.

Märkle, H., Saur, I.M.L., and Stam, R. (2022) Evolution of resistance (R) gene specificity. Essays in Biochemistry, 66, 551–560.

Martin, R., Qi, T., Zhang, H., Liu, F., King, M., Toth, C., et al. (2020) Structure of the activated ROQ1 resistosome directly recognizing the pathogen effector XopQ. Science, 370, eabd9993.

Mirdita, M., Schütze, K., Moriwaki, Y., Heo, L., Ovchinnikov, S., and Steinegger, M. (2022) ColabFold: making protein folding accessible to all. Nat Methods, 19, 679–682.

Möller, M. and Stukenbrock, E.H. (2017) Evolution and genome architecture in fungal plant pathogens. Nat Rev Microbiol, 15, 756–771.

Ortiz, D., Chen, J., Outram, M.A., Saur, I.M.L., Upadhyaya, N.M., Mago, R., et al. (2022) The stem rust effector protein AvrSr50 escapes Sr50 recognition by a substitution in a single surface-exposed residue. New Phytologist, 234, 592–606.

Pennington, H.G., Jones, R., Kwon, S., Bonciani, G., Thieron, H., Chandler, T., et al. (2019) The fungal ribonuclease-like effector protein CSEP0064/BEC1054 represses plant immunity and interferes with degradation of host ribosomal RNA. PLoS Pathog, 15, e1007620.

Pettersen, E.F., Goddard, T.D., Huang, C.C., Meng, E.C., Couch, G.S., Croll, T.I., et al. (2021) UCSF CHIMERAX : Structure visualization for researchers, educators, and developers. Protein Science, 30, 70–82.

Pierce, B.G., Wiehe, K., Hwang, H., Kim, B.-H., Vreven, T., and Weng, Z. (2014) ZDOCK server: interactive docking prediction of protein–protein complexes and symmetric multimers. Bioinformatics, 30, 1771–1773.

Prigozhin, D.M. and Krasileva, K.V. (2021) Analysis of intraspecies diversity reveals a subset of highly variable plant immune receptors and predicts their binding sites. The Plant Cell, 33, 998–1015.

Prigozhin, D.M., Sutherland, C.A., Rangavajjhala, S., and Krasileva, K.V. (2024) Majority of the highly variable NLRs in maize share genomic location and contain additional target-binding domains. MPMI, MPMI-05-24-0047-FI.

Rueden, C.T., Schindelin, J., Hiner, M.C., DeZonia, B.E., Walter, A.E., Arena, E.T., and Eliceiri, K.W. (2017) ImageJ2: ImageJ for the next generation of scientific image data. BMC Bioinformatics, 18, 529.

Sánchez-Vallet, A., Fouché, S., Fudal, I., Hartmann, F.E., Soyer, J.L., Tellier, A., and Croll, D. (2018) The Genome Biology of Effector Gene Evolution in Filamentous Plant Pathogens. Annu. Rev. Phytopathol., 56, 21–40.

Sarris, P.F., Cevik, V., Dagdas, G., Jones, J.D.G., and Krasileva, K.V. (2016) Comparative analysis of plant immune receptor architectures uncovers host proteins likely targeted by pathogens. BMC Biol, 14, 8.

Saur, I.M., Bauer, S., Kracher, B., Lu, X., Franzeskakis, L., Müller, M.C., et al. (2019) Multiple pairs of allelic MLA immune receptor-powdery mildew AVRA effectors argue for a direct recognition mechanism. eLife, 8, e44471.

Saur, I.M.L., Bauer, S., Lu, X., and Schulze-Lefert, P. (2019) A cell death assay in barley and wheat protoplasts for identification and validation of matching pathogen AVR effector and plant NLR immune receptors. Plant Methods, 15, 118.

Segretin, M.E., Pais, M., Franceschetti, M., Chaparro-Garcia, A., Bos, J.I.B., Banfield, M.J., and Kamoun, S. (2014) Single Amino Acid Mutations in the Potato Immune Receptor R3a Expand Response to Phytophthora Effectors. MPMI, 27, 624–637.

Seong, K. and Krasileva, K.V. (2021) Computational Structural Genomics Unravels Common Folds and Novel Families in the Secretome of Fungal Phytopathogen Magnaporthe oryzae. MPMI, 34, 1267–1280.

Seong, K. and Krasileva, K.V. (2023) Prediction of effector protein structures from fungal phytopathogens enables evolutionary analyses. Nat Microbiol, 8, 174–187.

Starr, T.N. and Thornton, J.W. (2016) Epistasis in protein evolution. Protein Science, 25, 1204–1218.

Tamborski, J., Seong, K., Liu, F., Staskawicz, B.J., and Krasileva, K.V. (2023) Altering Specificity and Autoactivity of Plant Immune Receptors Sr33 and Sr50 Via a Rational Engineering Approach. MPMI, 36, 434–446. The PyMOL Molecular Graphics System.

Wee, J. and Wei, G.-W. (2024) Evaluation of AlphaFold 3’s Protein–Protein Complexes for Predicting Binding Free Energy Changes upon Mutation. J. Chem. Inf. Model., 64, 6676–6683.

Yan, Y., Tao, H., He, J., and Huang, S.-Y. (2020) The HDOCK server for integrated protein–protein docking. Nat Protoc, 15, 1829–1852.

Zdrzałek, R., Stone, C., De La Concepcion, J.C., Banfield, M.J., and Bentham, A.R. (2023) Pathways to engineering plant intracellular NLR immune receptors. Current Opinion in Plant Biology, 74, 102380.

Zdrzałek, R., Xi, Y., Langner, T., Bentham, A.R., Petit-Houdenot, Y., De La Concepcion, J.C., et al. (2024) Bioengineering a plant NLR immune receptor with a robust binding interface toward a conserved fungal pathogen effector. Proc. Natl. Acad. Sci. U.S.A., 121, e2402872121.

Zhao, Y.-B., Liu, M.-X., Chen, T.-T., Ma, X., Li, Z.-K., Zheng, Z., et al. (2022) Pathogen effector AvrSr35 triggers Sr35 resistosome assembly via a direct recognition mechanism. Sci. Adv., 8, eabq5108.

