## Supplemental Figures for "Resurrection of the Plant Immune Receptor Sr50 to Overcome Pathogen Immune Evasion"

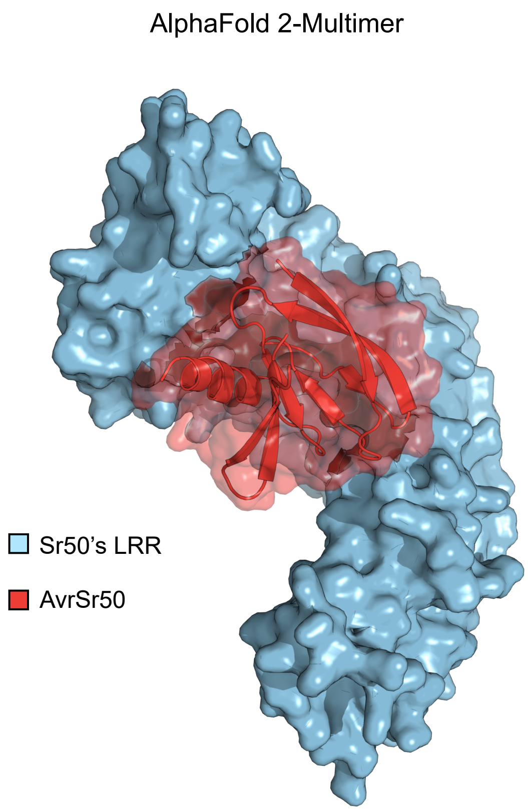


**Figure S1. The predicted complex structure of Sr50 and AvrSr50 by AlphaFold 2 and 3**

The full-length sequences of Sr50 and the mature protein sequences of AvrSr50 were used to predict the multimer structure with ColabFold and AlphaFold-Multimer (left) as well as AlphaFold 3 (right). The best prediction showed the ipTM score of 0.55 and 0.33, respectively. The leucine-rich repeat (LRR) domain of Sr50 and AvrSr50 are shown. Some parts of a loop between β2–β3, including an unstructured region, is removed from AvrSr50 for visualization (positions 42-66). AlphaFold 3 failed to predict the structure of AvrSr50, as indicated by the absence of its alpha helix.

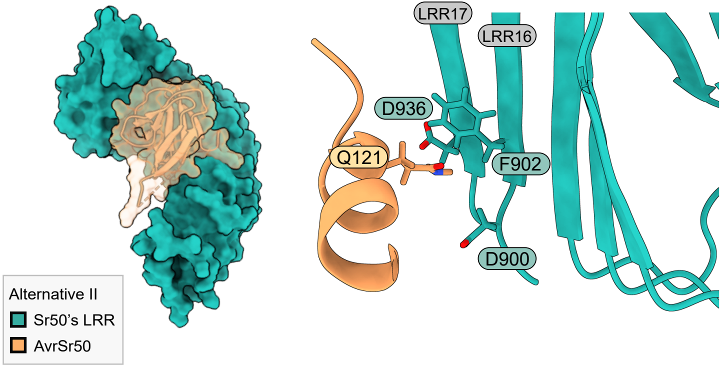


**Figure S2. Alternative structural hypotheses derived from molecular docking simulations**

The local environment around AvrSr50's Q121 is visualized in Alternative model II. This model was excluded from our initial screening, as the model did not align with our simplified assumption.

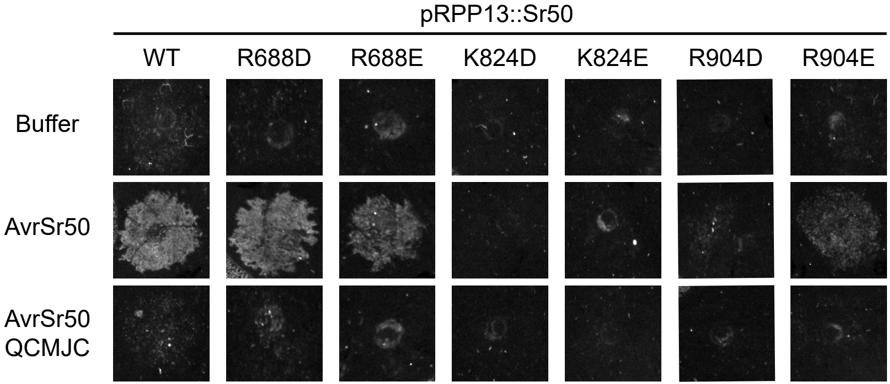


**Figure S3. The tested Sr50 mutants do not lead to AvrSr50$^{QCMJC}$-dependent cell death**

The cell death phenotypes on *Nicotiana benthamiana* leaves at three days post infiltration. *Agrobacterium* carrying wild type *Sr50* (WT) or a *Sr50* with an indicated substitution was co-infiltrated with buffer or *Agrobacterium* transformed to express AvrSr50 or AvrSr50$^{QCMJC}$. The optical density ($OD_{600}$) was set to 0.3 for receptors and effectors, respectively.

Residue index

**Mature protein** **Full-length protein**

**1**, **23**

AvrSr50 ARSLVKIDWSGSEYTILGAN
AvrSr50$^{QCMJC}$ ARSLIKTDWSGSEYTILGAN

**21**, **43**

AvrSr50 HYEEPNTGAAAQFPGTMTVD
AvrSr50$^{QCMJC}$ HYEEPNTGAAAQFPGTMAED

**41**, **63**

AvrSr50 DGRSPYIVRKLRNSSGKRFY
AvrSr50$^{QCMJC}$ DGRSPYIVRKLRNSSGKRFY

**61**, **83**

AvrSr50 VFTGHPQQPIVWNPHEEIEI
AvrSr50$^{QCMJC}$ VFTDHPQQPIIWNPHEEIEI

**81**, **103**

AvrSr50 QFNRKFLIAVLTEFEADSQV
AvrSr50$^{QCMJC}$ QFSRKYLIAVLTEFEADSKV

**101**, **123**

AvrSr50 FNHFARRQHR
AvrSr50$^{QCMJC}$ FTHFARRQHR

**Figure S4. Sequence variations between AvrSr50 and AvrSr50$^{QCMJC}$**

The pairwise sequence alignment between AvrSr50 and AvrSr50$^{QCMJC}$ (PDB:7MQQ). The mature protein sequences were used for both effectors. Two residue indexes are given based on the mature protein (green) and the full-length protein (blue).

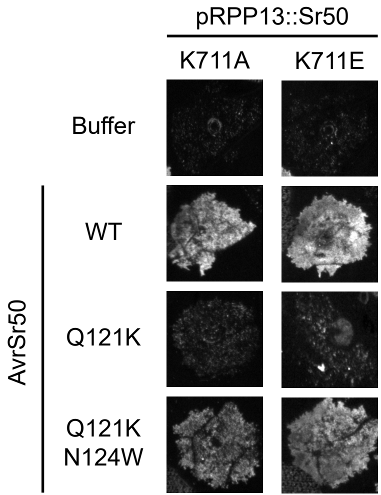


**Figure S5. The AvrSr50$^{Q121K/N124W}$ mutant increases cell death towards some Sr50 K711 mutants**

The cell death phenotypes at three days post infiltration. *Agrobacterium* carrying Sr50$^{K711A}$ or Sr50$^{K711E}$ was co-infiltrated with buffer or *Agrobacterium* with AvrSr50 variants.

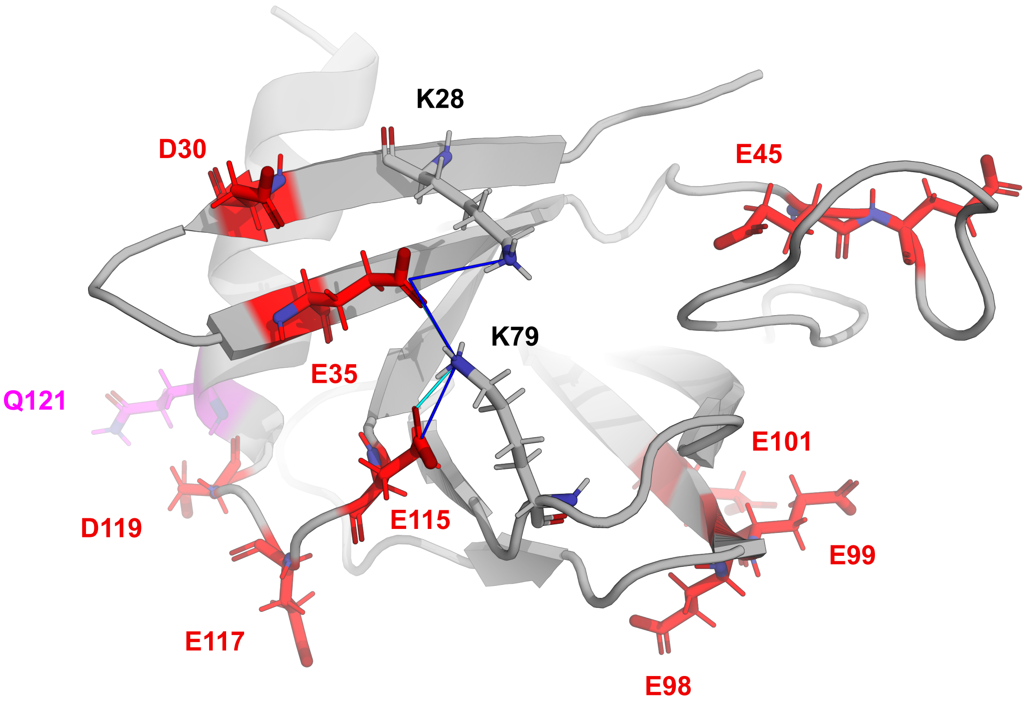


**Figure S6. AvrSr50's E35 and E115 may participate in protein stabilization**

The predicted structure of AvrSr50 with negatively charged amino acids visualized in red, Q121 in pink and two lysine residues in grey. The residue index is based on the full-length protein. The analysis with RING suggested that E35 and E115 may form ionic bonds (blue) and hydrogen bonds (sky blue) with K28 and K79 for protein stability.

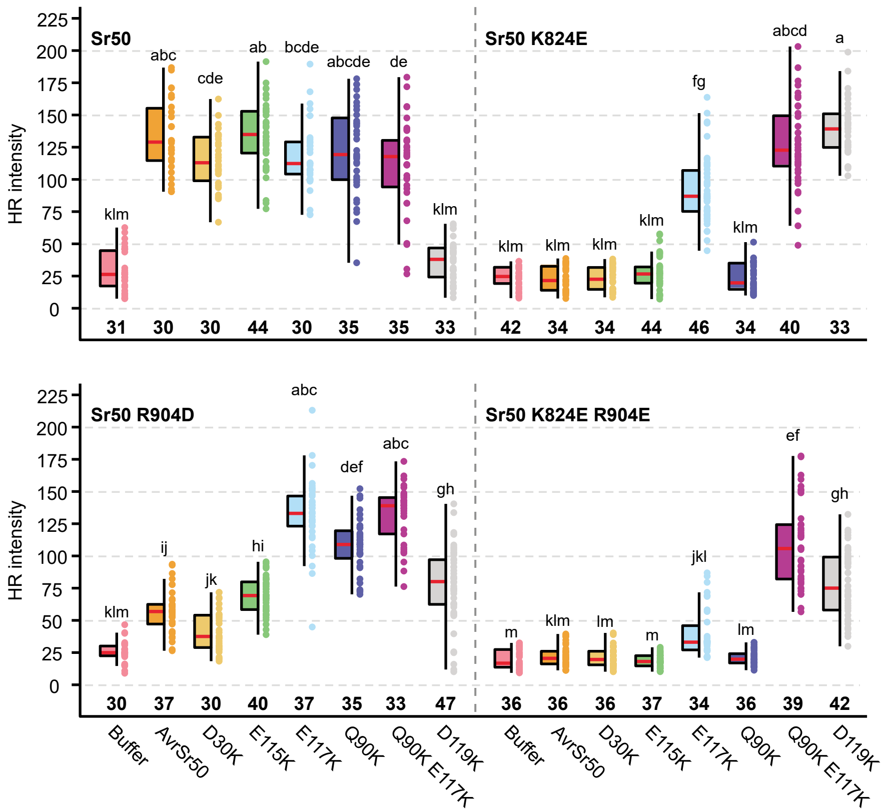


**Figure S7. Cell death phenotypes on *Nicotiana benthamiana***

Indicated pairs of receptors and effectors were co-infiltrated, and the hypersensitive response (HR) intensity was quantified at three days post infiltration. The statistical significance was accessed with one-way ANOVA followed by a post-hoc Tukey Honestly Significant Difference test. Groups sharing the same letter are not significantly different (P > 0.05). The numbers in bold indicate replicates.

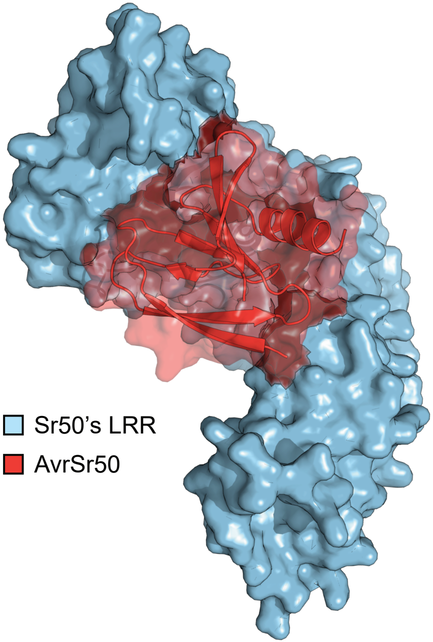


**Figure S8. A ColabDock model**

A model derived from ColabDock with the following constraints of the receptor and effector residues: K824 and E117, as well as R904 and E117.

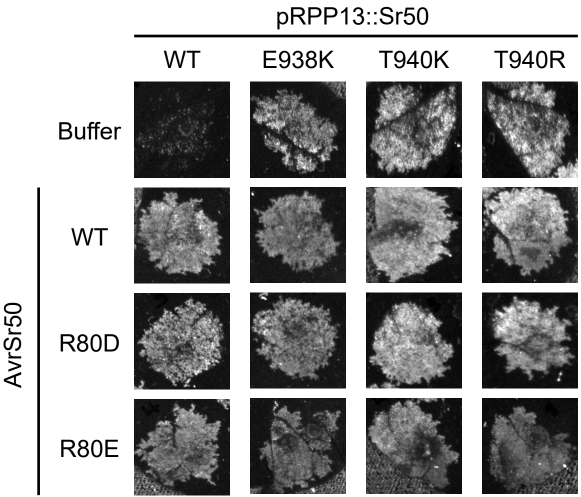


**Figure S9. Positively charged residues at positions 938 and 940 of Sr50 lead to severe auto-activity**

Cell death phenotypes on *Nicotiana benthamiana*. Indicated receptor and effector pairs were co-infiltrated, and the phenotypes were recorded at three days post infiltration. Representative HR images are displayed.

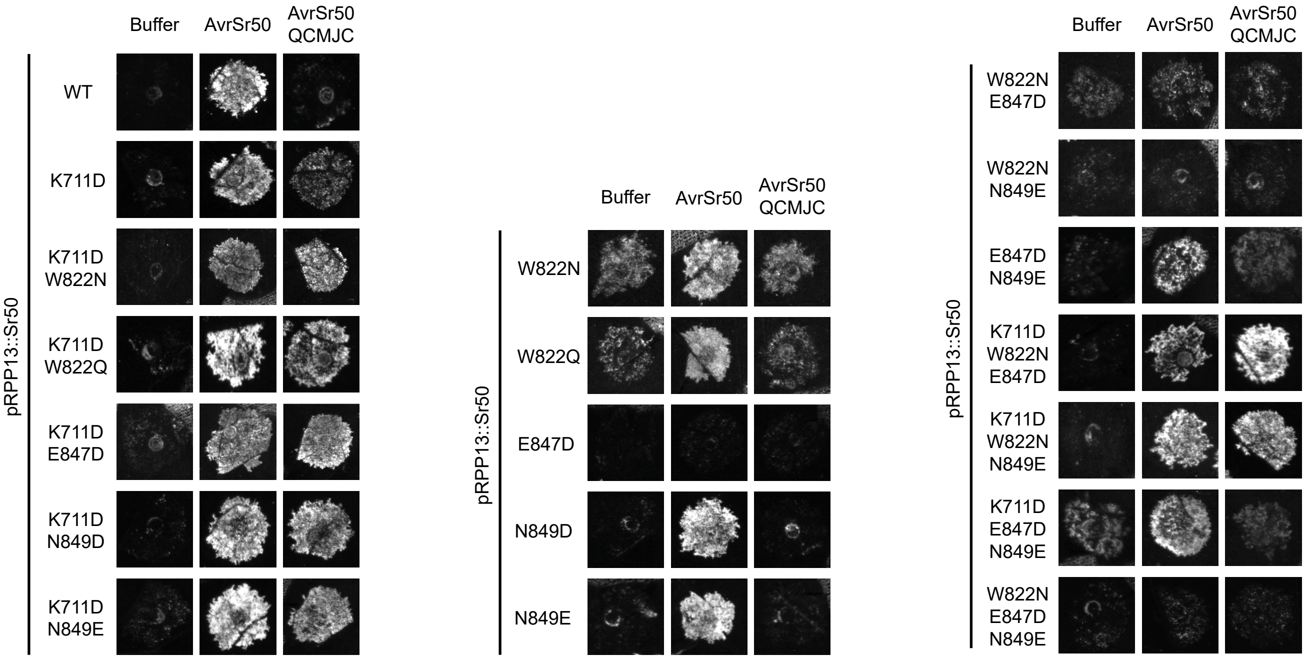


**Figure S10. Cell death phenotypes on *Nicotiana benthamiana***

Indicated pairs of receptors and effectors were co-infiltrated, and the hypersensitive response (HR) intensity was quantified at three days post infiltration. Representative HR images corresponding to the median intensity are displayed. **(b)** The statistical significance was accessed with one-way ANOVA followed by a post-hoc Tukey Honestly Significant Difference test. Groups sharing the same letter are not significantly different ($P > 0.05$). The numbers in bold indicate replicates.

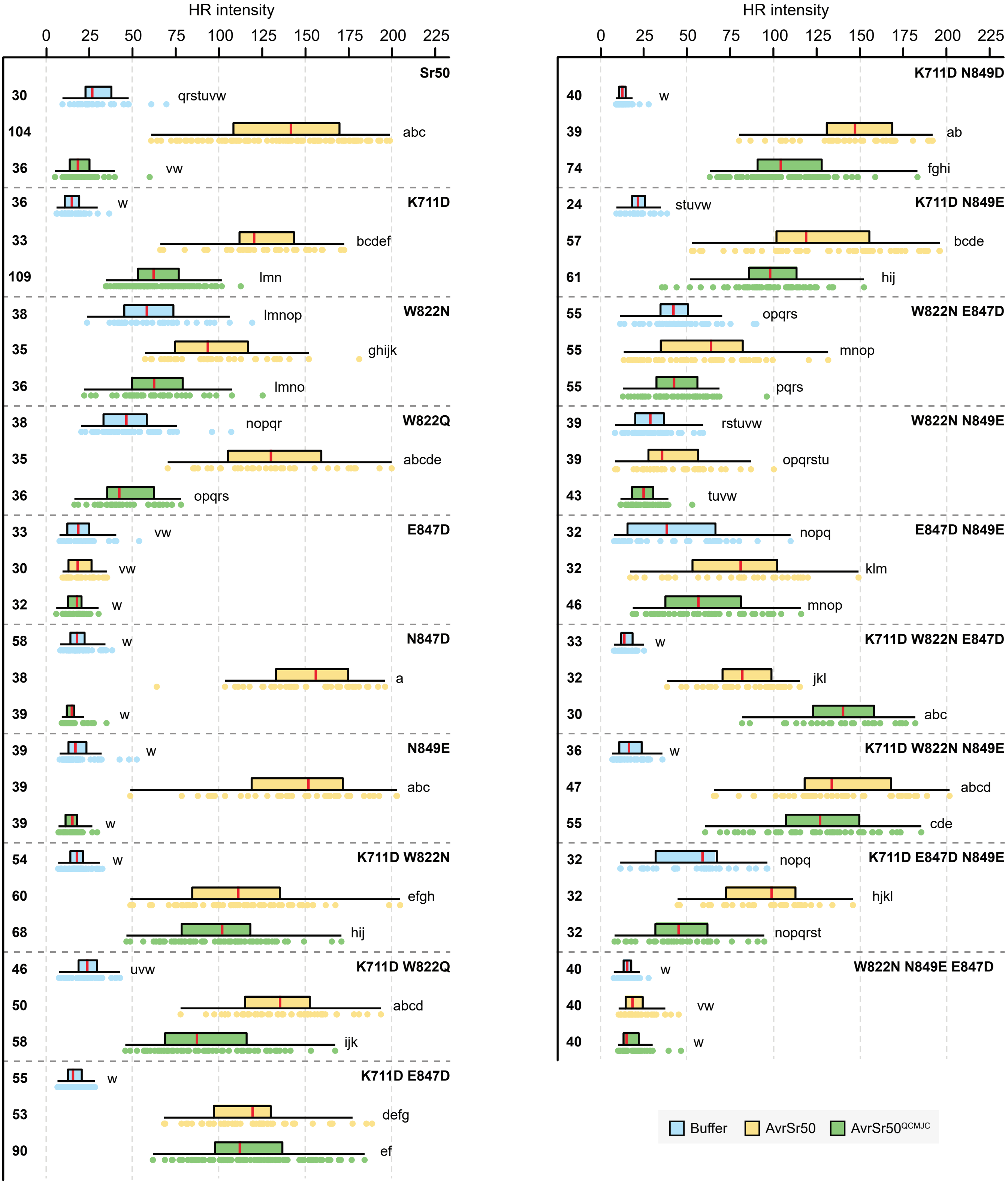


**Figure S11. Quantification of cell deaths for the engineered receptor mutants**

Indicated pairs of receptors and effectors were co-infiltrated, and the hypersensitive response (HR) intensity was quantified at three days post infiltration. The statistical significance was accessed with one-way ANOVA followed by a post-hoc Tukey Honestly Significant Difference test. Groups sharing the same letter are not significantly different (P > 0.05). The numbers in bold indicate replicates.

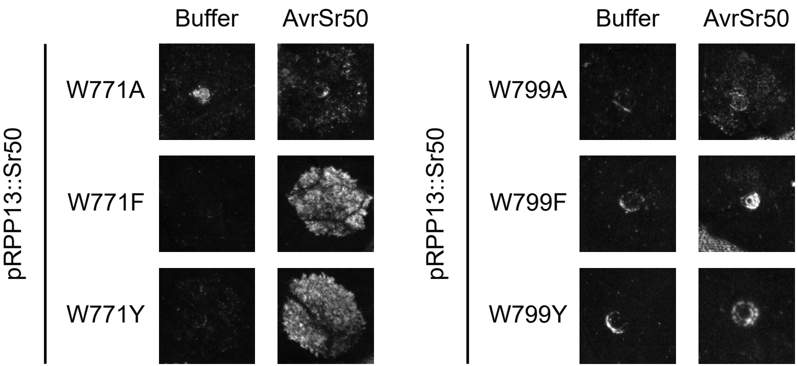


**Figure S12. Sr50's W799 is essential for AvrSr50 recognition**

The cell death phenotypes at three days post infiltration. *Agrobacterium* carrying Sr50 with an indicated mutation was co-infiltrated with infiltration buffer or *Agrobacterium* with AvrS50.

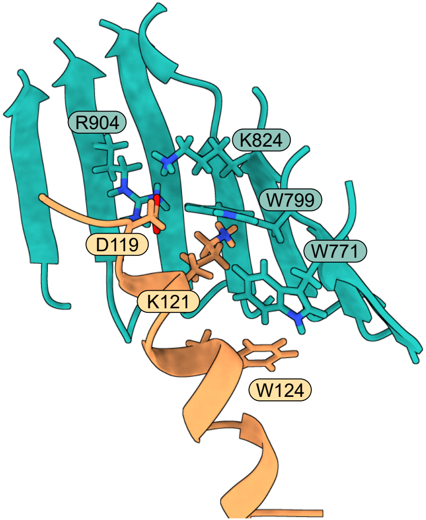


**Figure S13. AvrSr50's N124 is proximal to Sr50's W771**

Predicted model of Sr50$^{K711D}$ and AvrSr50$^{Q121K/N124W}$ with an ipTM score of 0.803. The local environment around K121 is visualized.

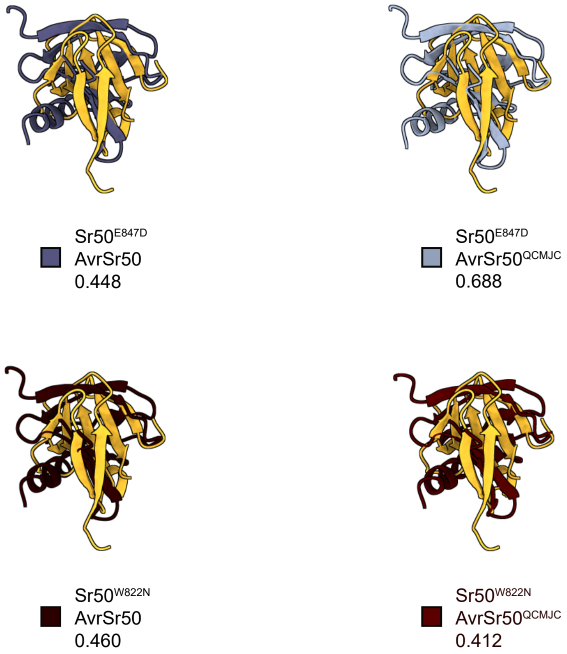


**Figure S14. The E847D and W822N substitutions do not change the behavior of AlphaFold**

Structural superposition of AvrSr50 in AlphaFold 2 (AF2) models compared to Model IV. Predicted protein complex structures were superposed against Model IV to keep receptor coordinates consistent. The AvrSr50 structures were then visualized. The yellow structure is AvrSr50 from Model IV. Labels indicate the receptors and effectors used for prediction and their associated confidence scores (ipTM).
